# Altered costimulatory signals and hypoxia support chromatin landscapes limiting the functional potential of exhausted T cells in cancer

**DOI:** 10.1101/2021.07.11.451947

**Authors:** B. Rhodes Ford, Natalie L. Rittenhouse, Nicole E. Scharping, Paolo D. A. Vignali, Andrew T. Frisch, Ronal Peralta, Greg M. Delgoffe, Amanda C. Poholek

## Abstract

Immunotherapy has changed cancer treatment with major clinical successes, but response rates remain low due in part to elevated prevalence of dysfunctional, terminally exhausted T cells. However, the mechanisms promoting progression to terminal exhaustion remain undefined. We profiled the histone modification landscape of tumor-infiltrating CD8 T cells throughout differentiation, finding terminally exhausted T cells possessed chromatin features limiting their transcriptional potential. Active enhancers enriched for bZIP/AP-1 transcription factor motifs lacked correlated gene expression, which were restored by immunotherapeutic costimulatory signaling. Epigenetic repression was also driven by an increase in histone bivalency, which we linked directly to hypoxia exposure. Our study is the first to profile the precise epigenetic changes during intratumoral differentiation to exhaustion, highlighting their altered function is driven by both improper costimulatory signals and environmental factors. These data suggest even terminally exhausted T cells remain poised for transcription in settings of increased costimulatory signaling and reduced hypoxia.

## INTRODUCTION

Immunotherapeutic approaches in cancer aim to stimulate, reinvigorate, or augment a T cell response to cancer cells, driven primarily by CD8 T cells. Most notable among these is antibody-mediated blockade of ‘checkpoint’ molecules like CTLA-4 and PD-1(*1*). CD8 T cells ultimately fail in cancer due to their differentiation into an exhausted state, leading to limited proliferative capacity, metabolic dysfunction, lower cytokine expression and decreased killing capacity(*2, 3*). The goal of PD-1 blockade, in principle, is to block inhibitory signaling on effector or exhausted T cells within the tumor. However, it is now clear that PD-1 blockade acts on less differentiated T cells in the tumor, inducing their acquisition of a cytotoxic phenotype, rather than direct PD-1 blockade on exhausted T cells(*4, 5*). In fact, the prevalence of exhausted T cells in tumors has been shown to correlate with resistance, rather than response, to PD-1 blockade(*6, 7*). Thus, it is critical to understand how T cells progress to terminal exhaustion and what underlies their dysfunction to design strategies that would specifically target these tumor-specific T cells and restore adequate function.

While exhaustion may have previously been thought of as a functional state of T cells, it is now well established that the transcriptome and epigenome of exhausted T cells are independent states from effector and memory T cells indicating the permanence of the exhausted cellular differentiation lineage(*6, 8, 9*). This differentiation is driven by the convergence of various chronic signals, including those from the antigen receptor as well as metabolic stress responses. Differentiation to exhaustion, however, is a progressive process, and progenitor or stem-like states have been identified that are transcriptionally distinct from terminally exhausted cells. Progenitor cells have increased proliferative potential, effector cytokine production, metabolic capacity and can be identified by reduced expression of inhibitory receptor expression. Transcription factor expression differences are also associated with subsets of exhaustion, with TCF-1 expression highest in progenitor cells while Tox expression increases in terminal cells(*10–14*). A recent study demonstrated that these states also had distinct chromatin landscapes as measured by ATAC-seq, which can interrogate open regions of chromatin(*6*).

The epigenetic states controlling transcription are complex. While the full understanding of how histone modifications contribute to gene regulation is still an active area of research, several well described modifications have been associated with specific chromatin states, which can provide a more nuanced view of epigenomic relationships that regulate the transcriptome of independent cellular states(*15, 16*). Recent advances in low cell input chromatin assays have overcome technical hurdles in assessing the epigenetic landscape as defined by histone modification, rather than solely accessibility, from tumor-infiltrating immune cells. Using Cleavage Under Targets & Release Under Nuclease (CUT&RUN), we set out to determine the epigenetic landscape associated with the progression to terminal exhaustion in tumor-infiltrating T cells by interrogating 4 well-described histone modifications(*17*). In addition, we determined the relationship of Tox, a transcription factor associated with terminal exhaustion with the epigenetic landscape and transcriptome. Here, we describe unexpected chromatin states in terminally exhausted CD8^+^ T cells: an active chromatin landscape where a significant fraction of genes does not correlate with active transcription, and an increased fraction of chromatin that exhibits bivalency, typically indicative of a stem cell like state. In this study, we describe the relationship of the epigenome with the transcriptome and find that impaired costimulatory signaling and exposure to hypoxia in the tumor microenvironment contribute to altered gene regulation identified via the epigenome to promote exhaustion. Our study for the first time explores in depth the epigenome of T cell exhaustion and lays the groundwork for future exploration of how the dysregulated epigenome of exhausted T cells may be targeted to further repurpose these cells therapeutically.

## RESULTS

### CUT&RUN reveals distinct chromatin features of progenitor and terminally exhausted tumor-infiltrating T cells at steady state

Previous studies using scRNA-seq and ATAC-seq demonstrated that progenitor and terminally exhausted states have distinct transcriptomes and chromatin accessibility in both chronic viral infection and tumors, yet how specific epigenetic changes contribute to the transition from progenitor to terminal exhaustion has not been explored(*6, 8, 9, 18, 19*). We employed B16 melanoma, an aggressive murine tumor that predictably promotes T cell exhaustion and is insensitive to PD-1 blockade monotherapy(*20, 21*) to better understand the relationship between the epigenome and the transcriptome of CD8^+^ tumor-infiltrating lymphocytes (TIL) as they progressively differentiated from a progenitor state (PD-1^lo^ Tim3^-^) to a terminally exhausted state (PD-1^hi^ Tim3^+^). Endogenous CD8^+^ T cells were isolated from day 14 B16 tumors and flow sorted into 4 populations based on expression of PD-1 and Tim3 (PD-1^lo^, PD-1^mid^, PD-1^hi^, PD-1^hi^ Tim3^+^) for analysis by RNA-seq to assess transcriptomes (Fig 1A). As naïve T cells have extremely condensed chromatin, we sorted antigen-experienced CD8 T cells (CD44^hi^) from paired draining LNs (dLNs) and compared to TIL as a control(*22*). Transcriptome analysis of sorted populations confirmed all TIL subsets segregated away from activated CD44^hi^ CD8 T cells in the dLN (Fig. 1B). Further, PD-1^lo^ and PD-1^mid^ TIL were closely related, while PD-1^hi^ and PD-1^hi^ Tim3^+^ cells were similar to each other suggesting a transition between PD-1^mid^ to PD-1^hi^ cells that discriminated the progenitor exhausted and terminal exhausted states (Fig 1B).

**Figure 1.**
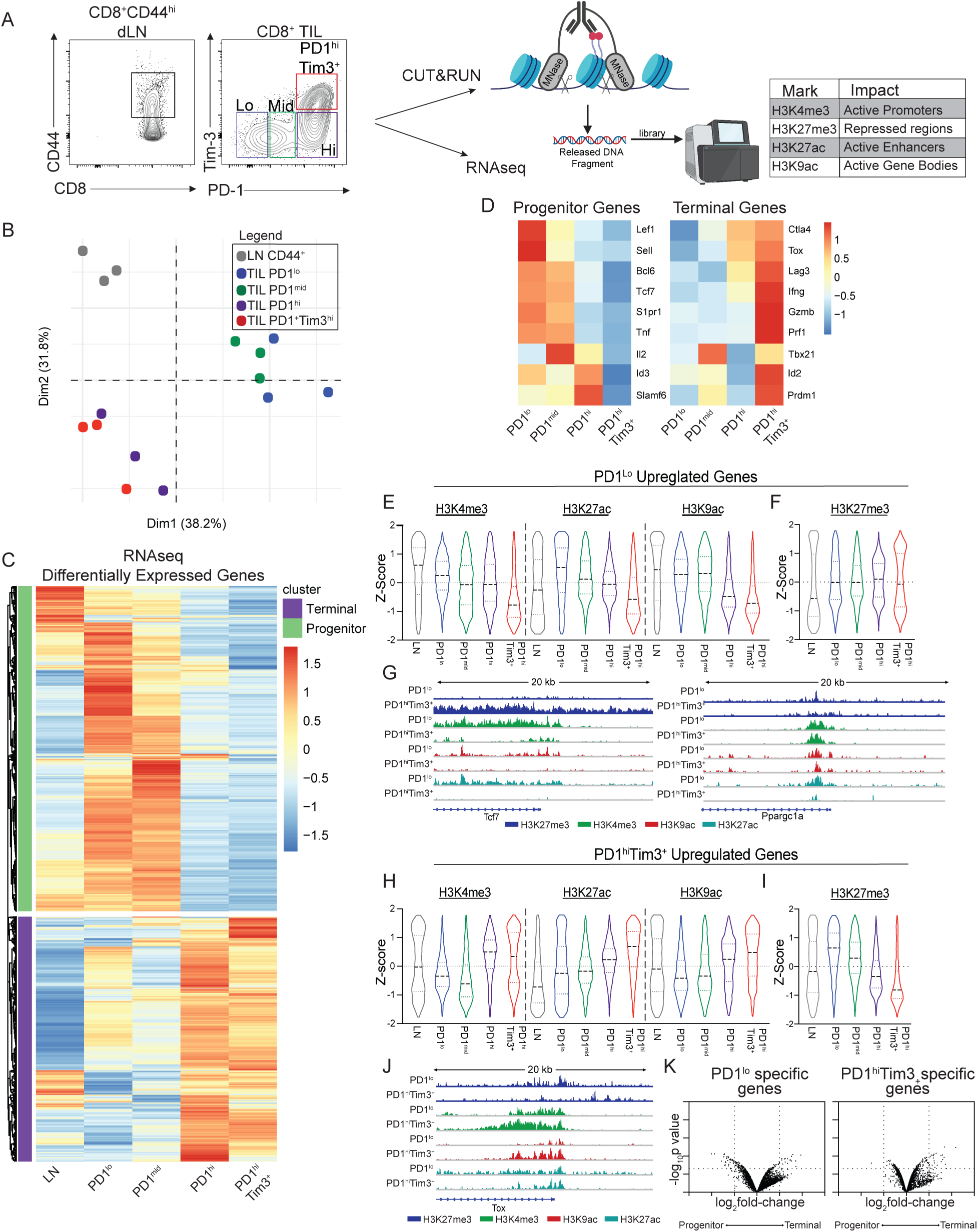
Transcriptional signatures of progenitor and terminally exhausted cells are set by histone landscapes. (A) Sort strategy and schematic of RNAseq and CUT&RUN of CD44^hi^ CD8 T cells from the draining LN and B16 melanoma tumors. TIL were sorted by PD-1 and Tim-3 expression into indicated subsets. (B) PCA of transcriptomes of LN CD44^hi^ CD8 T cells and TIL subsets. (C) Heatmap shows DESeq2 of log2 normalized transcript expression of differentially expressed genes between LN CD44^hi^ CD8 T cells and TIL subsets. Values are transformed log2(TPM). (D) Heatmaps of select differentially expressed genes known to be associated with progenitor and terminally exhausted T cell programs. (E and F) Violin plots of H3K4me3, H3K27ac, H3K9ac (E), and H3K27me3 (F) coverage of genes upregulated in PD-1^lo^ cells. Z-score indicates tag counts per million of 10kb regions surrounding the TSS. (G) IGV plots of H3K27me3, H3K4me3, H3K9ac, and H3K27ac at *Tcf7* and *Ppargc1a* in indicated subsets. (H and I) Violin plots of H3K4me3, H3K27ac, H3K9ac (H), and H3K27me3 (I) coverage of genes upregulated in PD-1^hi^ Tim3^+^ cells. Z-score indicates tag counts per million of 10kb regions surrounding the TSS. (J) IGV plots of H3K27me3, H3K4me3, H3K9ac, and H3K27ac at *Tox*. (K) Volcano plot of fold change and p value of changes in chromatin accessibility determined by ATAC-seq (GSE122713) in progenitor and terminally exhausted cells from B16 melanoma at genes specific to PD-1^lo^ and PD-1^hi^Tim3^+^ CD8 TIL (Log2 fold-change > 1 and log10 p value < 0.05). Coverage was determined for 10kb regions surrounding the TSS of each gene. RNAseq data generated from three individual mice treated as biological replicates. CUT&RUN for each mark was performed twice with 8-10 mice pooled per experiment before sorting.

Analysis of differentially expressed genes (DEG) confirmed that PD-1^lo^ and PD-1^mid^ cells shared a transcriptional profile, while PD-1^hi^ and PD-1^hi^ Tim3^+^ cells were similar (Fig 1C). DEG in PD-1^lo^ cells confirmed a gene signature similar to that previously described for progenitor exhausted cells while PD-1^hi^ Tim3^+^ cells had a signature consistent with terminal exhaustion, such as increased expression of *Slamf6*, *Tcf7*, *Bcl6*, *Id3*, *Lef1*, and *Tnf* in progenitor cells while genes such as *Lag3*, *Tox*, *Ifng*, *Prdm1* and *Id2* were upregulated in PD-1^hi^ Tim3^+^ cells (Fig 1D). Furthermore, Pearson correlation with previously published RNAseq of progenitor and terminal cells isolated from tumors demonstrated that PD-1^lo^ cells were more similar to Slamf6^+^ progenitor cells, while PD-1^hi^ Tim3^+^ were more similar to Slamf6^-^ Tim3^+^ terminally exhausted cells(*6*) (Fig S1A). Thus, gating cells via PD-1 and Tim3 expression sufficiently identified both progenitor and terminal states of exhaustion that likely transition as cells move from the PD-1^mid^ to the PD-1^hi^ stage.

Next, we used CUT&RUN to generate a foundational epigenetic map of chromatin states to assess active chromatin (H3K4me3, H3K9ac and H3K27ac) and repressed chromatin (H3K27me3) as TIL progressively differentiated (Fig 1A). To determine if the epigenetic states that underlie the transcriptomes of progressive exhaustion were consistent with gene expression, 20kb regions of chromatin around the TSS of DEG upregulated in PD-1^lo^ progenitor cells or PD-1^hi^ Tim3^+^ terminal cells were assessed. Indeed, genes upregulated in PD-1^lo^ cells displayed an increased amount of active chromatin (H3K4me3, H3K27ac, H3K9ac) compared to terminally exhausted cells (PD-1^hi^ Tim3^+^) and had less of the repressive mark H3K27me3 (Fig 1E,F). Interestingly terminally exhausted cells (PD-1^hi^ Tim3^+^) exhibited a bimodal distribution of H3K27me3 at progenitor-specific genes (Fig 1F). In terminal cells, some progenitor-specific genes had increased H3K27me3 (i.e., *Tcf7*), indicative of chromatin repression as cells differentiated to a terminal state. Yet some genes remained low for H3K27me3 (i.e. *Ppargc1a*), suggesting the loss of active chromatin may also be sufficient to downregulate gene expression in terminal cells (Fig 1F,G). Thus, for progenitor-specific genes to become downregulated in terminally exhausted cells, changes to the epigenome occur that drive either decreases in active chromatin or increases in repressive chromatin.

A similar pattern was observed for genes specific to terminal exhaustion with increased active chromatin (H3K4me3, H3K27ac, H3K9ac) at genes upregulated in PD-1^hi^ Tim3^+^ compared to PD-1^lo^ cells (Fig 1H). In contrast to the bimodal distribution of H3K27me3 for progenitor-specific genes, H3K27me3 at terminal specific genes was significantly increased in PD-1^lo^ cells, and dramatically reduced as cells progress to PD-1^hi^ Tim3^+^. These data suggest the progression to terminal exhaustion requires H3K27 demethylation for genes to become upregulated in terminally exhausted cells (Fig 1I). For example, at the *Tox* locus, H3K27me3 is reduced in PD-1^hi^ Tim3^+^ cells, while active marks are increased (Fig 1J). We next determined if changes in chromatin accessibility measured by ATAC-seq in publicly available datasets had similar alterations as histone modifications when looking at DEGs in progenitor and terminal states of exhaustion(*6*). Surprisingly, we found limited changes in accessibility suggesting changes in histone modifications provides a more nuanced view of epigenomic control of transcriptional programs for states of exhaustion (Fig 1K). Taken together, these data suggest that gene expression changes as progenitor cells progress to exhaustion are due to changes in the epigenome that regulate transcriptional control of gene expression.

We next compared the B16 TIL subsets to “bona fide” effector CD8 T cells generated using antigen-specific day 8 effector CD8 T cells (OT-I T cells) responding to an acute viral infection, *Vaccinia* virus expressing ovalbumin (*Vaccinia*-OVA) (Fig S1B). TIL segregated away from effector cells while progenitor and terminally exhausted T cells separated from each other in a manner similar to analysis of activated dLN cells (Fig S1C). DEG analysis identified transcriptional signatures unique to *Vaccinia*-OVA effector T cells, TIL progenitor, or TIL terminal exhaustion states, suggesting effector cells are transcriptionally distinct from both TIL exhaustion states. However, some genes which were highly expressed by effector cells had low to moderate expression in progenitor cells compared to terminal (Fig S1D). Chromatin states in effector cells tracked with gene expression, i.e. effector genes had increased active chromatin (H3K4me3) and decreased repressive chromatin (H3K27me3) (Fig S1E). Interestingly, both H3K4me3 and H3K27me3 were increased in terminally exhausted cells at effector gene loci (Fig S1E). The epigenetic state of effector cells at progenitor- and terminal-specific genes also had a mixed profile. H3K4me3 was decreased at progenitor genes, with bimodal distribution at terminal specific genes, suggesting some genes associated with terminal exhaustion may have active chromatin in effector cells (Fig S1F,G). We further compared our CUT&RUN datasets in B16 TIL subsets to previously published ChIP-seq analysis of naïve, effector and memory CD8 T cells responding to Listeria OVA(*23*). In line with our above analysis in Vaccinia-OVA responding cells, H3K4me3 profiles in effector CD8 T cells had similarities to terminal exhaustion, while naïve, memory and memory precursor cells had distinct profiles of active promoters (Fig S1H). In contrast, active enhancers marked by H3K27ac showed little correlation between TIL subsets and effector, memory, or naïve cells (Fig S1I). Repressed chromatin, however, had similarities between memory and progenitor exhausted cells, suggesting a shared landscape of repressed genes (Fig S1J). Collectively, these data suggest states of exhaustion in TIL are distinct from naïve, effector and memory CD8^+^ T cells responding to infection at both the level of the transcriptome and epigenome. Thus, as CD8^+^ TIL undergo progressive differentiation to terminal exhaustion, changes in the transcriptional signatures are mediated by key histone modifications. Furthermore, active histone modifications correlate with shifts in the transcriptional programs from progenitor to terminal TIL in ways that are more nuanced than broad changes in chromatin accessibility.

### Terminally exhausted T cells are specifically characterized by regions of active chromatin with low correlative transcription

Histone modifications such as H3K4me3, H3K27ac and H3K9ac mark active chromatin permissive for gene transcription by promoting open chromatin and mediating enhancer-promoter loops(*15*). To identify active enhancers specific to progenitor or terminal states, we performed differentially enriched peak (DEP) analysis of H3K27ac across TIL subsets (Fig 2A). This generated 3 clusters, with 2 dominant clusters of enhancers specific to progenitor cells (cluster 2) or terminally exhausted cells (cluster 1). Analysis of gene expression associated with enhancers in progenitor cells indicated the majority of active enhancers (∼85%) expectedly correlated with increased gene expression in progenitor cells that progressively decreased as cells differentiated to terminal exhaustion (Fig 2B,D). In contrast, in terminally exhausted T cells, only ∼55% of their specific active enhancers correlated with gene expression; an extensive fraction (∼45%) had anticorrelative gene expression (Fig 2C,E). While distal enhancers may account for a fraction of these, it is unlikely that the total fraction of active enhancers correlating with gene expression would be significantly lower in PD-1^hi^ Tim3^+^ cells than in PD-1^lo^ progenitor cells (Fig 2D,E). In addition, important genes known to be involved in effector T cell differentiation and function were among both the terminally exhausted correlated enhancer group (*Havcr2, Prdm1,* and *Ifng*), and anticorrelated enhancers (*Il10*, *Runx2*, and *Irf8*) (Fig. S2). These data suggested that terminally exhausted cells had a preponderance of active enhancers without correlated gene expression.

**Figure 2.**
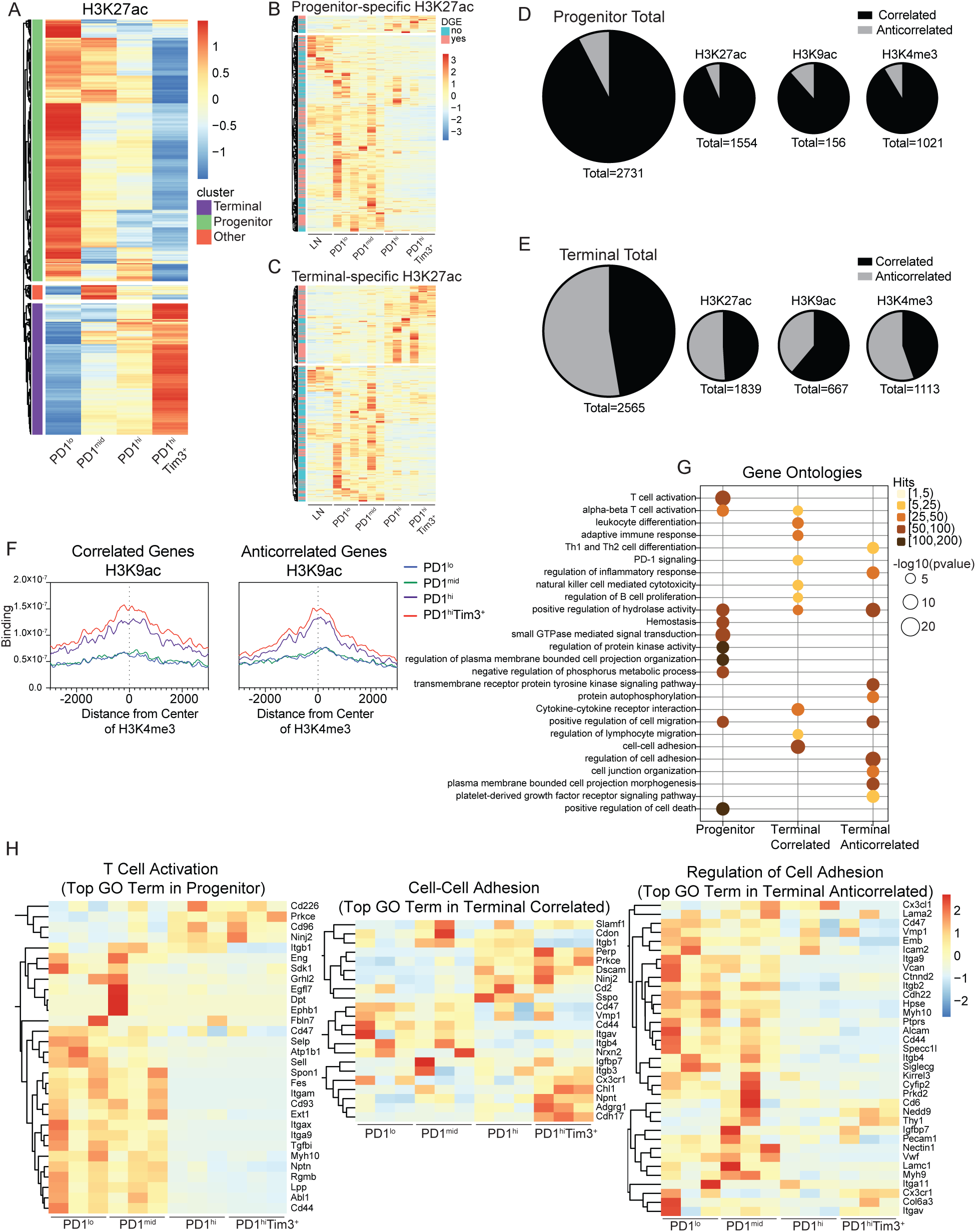
Active chromatin landscapes in terminally exhausted T cells do not correlate with gene expression. (A) Heatmap shows DESeq2 log2 normalized tag counts of H3K27ac at differential peaks identified between TIL subsets. (B and C) Heatmaps showing log2 normalized expression of genes nearest to progenitor-specific (B) and terminal-specific (C) H3K27ac peaks. Those genes that meet differentially expressed genes cutoffs (FC >2, p val < 0.05) marked in pink on the left side of heatmap. (D and E) Pie charts showing the number of genes with and without corresponding expression in progenitor (D) and terminally exhausted (E) T cells for H3K27ac, H3K9ac, H3K4me3, and the summed total of all three. (F) Histograms showing H3K4me3 coverage at correlated and anticorrelated H3K9ac peaks. (G) Bubble plot showing enrichment of gene ontology terms via Metascape in genes with active chromatin landscapes in indicated groups. (H) Heatmaps of log2 normalized expression of select genes enriched in GO pathways identified in (G).

When looking at the epigenetic profile of genes in both categories of terminally exhausted enhancer peaks, we noted that enhancer activity correlated with other marks of active chromatin (H3K9ac for active gene transcription, and H3K4me3 for active promoters) (Fig S2). We therefore explored how H3K9ac and H3K4me3 related to gene expression in progenitor or terminal populations. DEP analysis identified progenitor-specific and terminal-specific H3K4me3 and H3K9ac peaks which were further investigated for their relationship to gene transcription (Fig S3A,B). Similar to our analysis of active enhancers, progenitor-specific H3K4me3 and H3K9ac primarily correlated with gene expression, while terminal-specific H3K4me3 and H3K9ac identified a significant fraction of genes for which gene transcription was not correlated (Fig 2D,E, Fig S3C-F). In contrast, effector Vaccinia-OVA-elicited OT-I T cells primarily had correlative gene expression at active promoters suggesting the presence of a significant fraction of active chromatin with anticorrelative gene expression was unique to the terminal exhaustion state (Fig S3G,H).

We next determined the relationship between the three active chromatin marks profiled (H3K4me3, H3K9ac, H3K27ac) to determine if the combined presence of multiple active chromatin marks was key to drive gene expression. To test this, we asked how much H3K9ac and H3K27ac was present around H3K4me3-marked regions in the terminally exhausted correlated and anticorrelated gene groups. Surprisingly, we found similar amounts of H3K9ac and H3K27ac were associated with H3K4me3-specific peaks in both correlated genes and anticorrelated genes (Fig 2F, S3I). Taken together, these data indicate that, in general, the accumulation of active chromatin marks alone was insufficient to predict gene expression in terminally exhausted TIL.

We next explored the transcriptional pathways associated with progenitor-specific and terminal-specific active chromatin regions, breaking terminal-specific active chromatin into correlative and anticorrelative genes. GO pathway analysis demonstrated both overlapping and unique pathways to each group of genes, including T cell activation and lymphocyte or leukocyte pathways, PD-1 signaling, cell adhesion and cell migration (Fig 2G). Key genes involved in T cell activation and function were among those specifically expressed in progenitor- and terminal-specific regions of active chromatin (Fig 2H). Taken together, analysis of histone modifications by CUT&RUN revealed a number of genes critical for T cell activation and function that were not upregulated in exhausted T cells despite the presence of multiple active chromatin marks. Thus, terminally exhausted CD8 TIL have a unique chromatin landscape characterized by the presence of active histone modifications but without corresponding active gene transcription.

### The anticorrelated enhancers of terminally exhausted T cells are uniformly defined by AP-1 binding motifs

We next sought to understand why a significant fraction of genes in terminally exhausted CD8 T cells maintain active chromatin but lack gene expression. We hypothesized that increased H3K27me3 might lead to repression of these genes. However, H3K27me3 levels were similar at both correlated and anticorrelated genes, suggesting the presence of repressive chromatin marks did not explain why gene expression was reduced (Fig S3J). Active chromatin is permissive for transcription, but transcription factors are required to recruit the machinery to drive gene expression(*15*). We next asked which transcription factors were enriched within progenitor-specific and terminal-specific active chromatin. Progenitor-specific enhancers were enriched for motifs from the ETS family and TCF1, while terminal-specific enhancers had enrichment for the bZIP consensus sequence bound by AP-1 family members (Fig 3A). Terminal-specific H3K4me3 and H3K9ac peaks were also enriched for bZIP motifs, while progenitor-specific H3K4me3 and H3K9ac peaks had less specific enrichment of any motifs compared to terminal cells. We next explored if there were differences in motif enrichment in correlated vs anticorrelated genes within terminal-specific active chromatin marks. bZIPs were enriched in enhancers of both correlated genes and anticorrelated genes, however there were two notable motifs enriched only in correlated genes; Nur77 and the composite NFAT:AP-1 motif. (Fig 3B). Nur77 (encoded by *Nr4a1*) activity correlates with recent TCR stimulation and NFAT:AP-1 complexes occur in response to TCR signaling, suggesting TCR signals play driving roles in promoting gene expression in terminal exhaustion-specific genes(*19, 24, 25*). Taken together, these data suggest the presence of key transcription factors may regulate gene expression in terminally exhausted CD8 T cells where chromatin is already primed via the presence of active chromatin marks.

**Figure 3.**
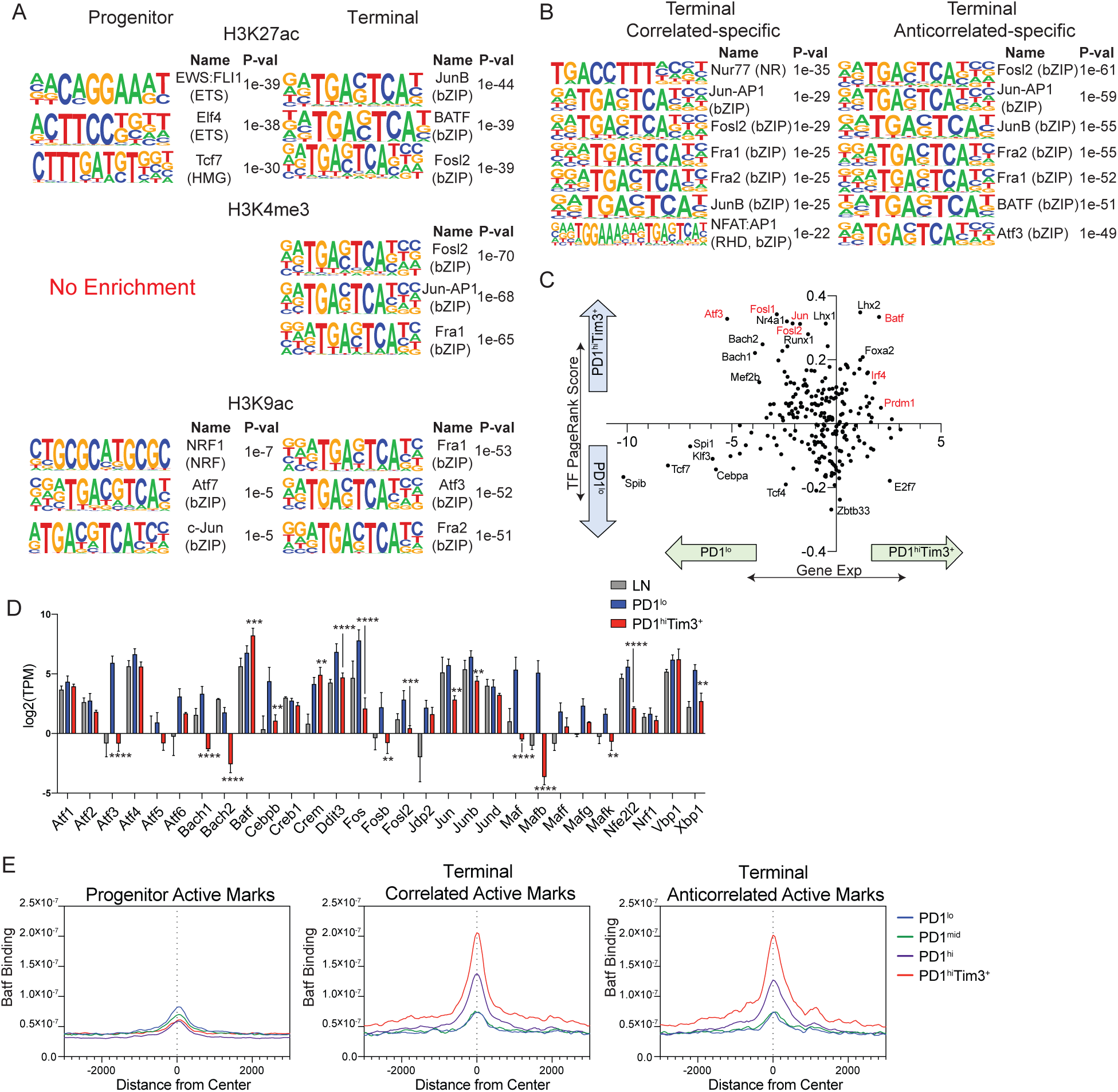
bZIP/AP-1 family motifs are enriched in active chromatin of terminally exhausted TIL. (A) HOMER motif analysis of DESeq2 defined peaks of H3K27ac, H3K4me3, and H3K9ac peaks identified in progenitor or terminally exhausted T cells. Terminally exhausted-specific peaks were used as background for progenitor exhausted-specific analysis, and vice versa. (B) HOMER motif analysis of DESeq2 defined correlated and anticorrelated peaks compiled for each mark. Progenitor exhausted-specific peaks were used as background for both correlated and anticorrelated analyses. (C) Dot plot showing fold change PageRank score (Y-axis) and gene expression (X-axis) for transcription factors. H3K4me3, H3K27ac, and gene expression data was provided to Taiji for PageRank analysis. Homer motif list was also provided to Taiji. (D) Gene expression of AP-1 family members in LN, PD-1^lo^, and PD-1^hi^Tim3^+^ (mean and SD). p value (DESeq2, generated by one-way ANOVA); **p<0.01,***p<0.001,****p<0.0001 (E) Histograms showing Batf coverage (n=1) at progenitor, correlated, and anticorrelated peaks active peaks defined by presence of H3K27ac, H3K9ac or H3K4me3.

Exhaustion is driven in part by chronic antigen stimulation, and our data suggested that altered TCR signaling may contribute to the loss of gene expression in terminally exhausted cells despite the presence of active chromatin. Chronic antigen stimulation driven exhaustion also occurs in the context of chronic viral infection, therefore we next sought to determine if bZIP motifs were enriched in terminally exhausted CD8 T cells responding to chronic viral infection. Using previously published ATACseq datasets (*6*) of progenitor and terminal CD8 T cells isolated from both chronic (clone 13) LCMV and B16 tumor models, we identified 5 clusters of peaks that could be distinguished as progenitor-specific (cluster 2), LCMV-specific (cluster 1), tumor-specific (cluster 3), LCMV-terminal specific (cluster 5) and tumor terminal-specific (cluster 4) (Fig S4A). Motif analysis identified Tcf7 and Lef1 consensus sequences in peaks specific to progenitor cells (cluster 2) (**S4B**). Like our analysis of active chromatin marks in TIL terminal cells, bZIP motifs were highly enriched in tumor-specific and tumor terminal-specific clusters (3 and 4). In contrast, ETS motifs were enriched in LCMV-specific and LCMV terminal-specific clusters (1 and 5) (Fig S4B). These data suggested that the enrichment of bZIP/AP-1 motifs in terminal active chromatin regions was specific to tumor-mediated exhaustion and not a general feature of all settings of T cell exhaustion such as chronic viral infection.

To further identify transcription factors that may play a key role in gene regulation in exhaustion subsets, we performed Taiji PageRank analysis which uses an integrative multi-omics data analysis framework to identify regulatory networks and candidate driver transcription factors(*23, 26*). Interestingly, highly ranked transcription factors in progenitor cells correlated with increased gene expression of those transcription factors (Fig 3C). In contrast, many of the transcription factors highly ranked in terminally exhausted cells had decreased gene expression, with notable exceptions including *Batf*, *Prdm1* and *Irf4* (Fig 3C). In fact, many of these downregulated but highly ranked transcription factors were AP-1 family members such as *Atf3*, *Jun*, and *Fosl2*. Based on the strong enrichment of bZIP transcription factors in terminal-specific active chromatin, we explored the expression of bZIP transcription factors in TIL subsets. We found that most bZIP transcription factors were downregulated in terminal cells, with few exceptions including Batf (Fig 3D). As a ChIP-seq-like technology, CUT&RUN is equivalently capable of measuring transcription factor binding, so we performed CUT&RUN for Batf in TIL. Although Batf was expressed in progenitor cells, it was not highly enriched at sites of active chromatin marks (H3K27ac, etc) in progenitor cells (Fig 3E). In contrast Batf was highly enriched at sites of active chromatin in terminally exhausted cells, however no differences were noted between correlated genes and anticorrelated genes. These data suggested that in line with the strong presence of bZIP motifs, the AP-1 family member Batf was bound specifically in terminal cells at places where active chromatin is present.

### Tox associates with key cell-specific transcription factors to promote exhaustion specific transcriptional programs

Tox is a transcription factor associated with terminal exhaustion that would not be identified by motif analysis as Tox does not bind a consensus motif but rather binds to secondary structure(*27*). To determine the relationship of Tox binding to the epigenetic changes identified above, we performed CUT&RUN for Tox in our TIL subsets. Although Tox increases in expression as cells progress towards terminal exhaustion, Tox is expressed at low levels in progenitor cells(*11, 12, 28-30*). Therefore, we were able to identify DEP of Tox in both progenitor and terminally exhausted CD8 T cells (Fig 4A). In progenitor cells, Tox bound several genes associated with the progenitor program, including *Tcf7*, *Bcl6*, *Bach2*, *Foxo1*, and *Id3*. While most Tox-bound genes in progenitor cells were associated with increased gene expression, several genes were decreased in progenitor cells, including *Il2ra* and *Irf4* (Fig 4B, S4C). Similarly, in terminal cells we found Tox bound to over 300 genes, many of which are associated with the transcriptional program of terminal cells, including *Havcr2* (encoding Tim3), *Tbx21*, *Ifng*, *Ctla4*, *Pdcd1*, *Tigit*, and *Tox* itself. Again, Tox binding was associated with both gene expression and repression (Fig 4B, 4C). Some notable Tox-bound genes that were associated with gene repression in terminal cells included *Irf8*, *Bcl11b* and *Il10*. Pathway analysis of Tox bound genes revealed transcriptional programs in both populations associated with T cell activation pathways, though enrichment was more significant in terminal cells than progenitor cells (Fig 4C). Motif analysis of Tox bound peaks revealed cell state-specific motifs; in progenitor cells, Tox binding was associated with Lef1 and Tcf7 motifs, while in terminal cells IRF:Batf motifs were strongly enriched (Fig 4D). IRF4, together with Batf has previously been associated with promoting T cell exhaustion. Indeed, we found Batf binding significantly overlapped Tox binding in terminal cells (Fig 4E). Furthermore IRF4 binding in activated CD8 T cells was more enriched at chromatin regions bound by Tox in terminally exhausted cells than progenitor cells, confirming enrichment of IRF4:Batf motifs of Tox bound peaks (Fig 4F). Next, we determined the relationship of Tox to active chromatin (K4me3, K27ac, and K9ac) in terminal cells described above. Tox binding overlapped with correlated and anticorrelated active mark peaks, though there was slightly more overlap in the correlated group (Fig 4G). Indeed, Tox was enriched at sites of active chromatin, and binding was slightly increased at sites that correlated with gene expression compared to anticorrelative genes (Fig 4H, Fig S4D). These data suggest Tox binds to active chromatin enriched in IRF:Batf composite motifs and works together with IRF4 and Batf to promote gene expression at key genes associated with the terminal exhaustion transcriptome. In contrast, in progenitor cells, Tox binds Tcf7 motifs to promote gene expression of the progenitor transcriptome.

**Figure 4.**
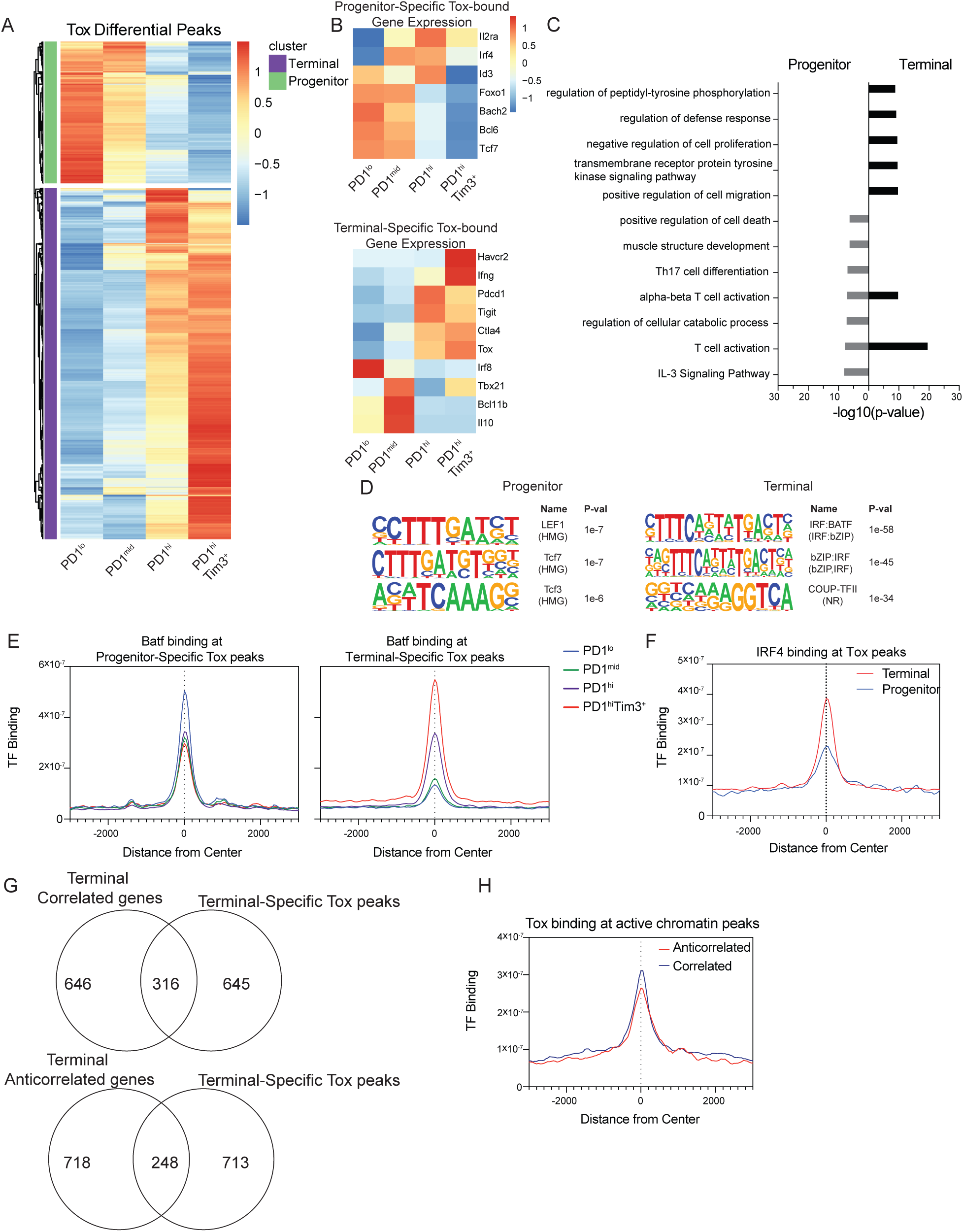
Tox associates with state-specific transcription factors TCF1 and Batf/IRF4. (A) Heatmap of log2 normalized DESeq2-defined differential Tox peaks in TIL subsets. (B) Heatmaps of log2 normalized selected genes associated with progenitor-specific and terminal-specific Tox peaks. (C) Bar graph showing -log10(p-value) values for GO term pathways identified by Metascape enriched for genes with progenitor and terminally exhausted-specific Tox binding. (D) HOMER motif analysis of DESeq2 defined peaks for progenitor and terminal-specific DESeq2-defined differential Tox peaks. Terminally exhausted-specific peaks were used as background for progenitor exhausted-specific analysis, and vice versa. (E) Histograms showing Batf coverage at progenitor and terminal-specific DESeq2-defined differential Tox peaks. n=1. (F) Histogram showing IRF4 binding (GSE54191) at progenitor and terminal-specific DESeq2-defined differential Tox peaks. IRF4 ChIP-seq is from *in vitro* activated T cells (*60*). (G) Venn diagrams comparing terminal specific peaks of active chromatin (K4me3, K27Ac, K9Ac) grouped by those with correlated or anticorrelated gene expression associated with terminal-specific Tox peaks (H) Histogram of Tox coverage in terminally exhausted cells at active chromatin defined as having anticorrelated and correlated gene expression as in Figure 2. CUT&RUN for Tox (n=2) and Batf (n=1) was performed with 8-10 mice pooled per experiment before sorting.

### Immunotherapies that alter costimulatory signaling drive gene expression of anticorrelated genes

Our study suggests that exhausted T cells, at the steady state, harbor enhancers defined by AP-1 binding sites that are not actively promoting gene transcription. AP-1 transcription factor activity is classically driven by costimulatory signaling cascades, and it is the lost costimulation that is thought to play a major role in driving exhaustion(*31*). NFAT activity in the absence of AP-1 can drive dysfunctional programs like anergy and exhaustion(*25*). Further, these pathways are primarily the targets of checkpoint blockade immunotherapies: PD-1 blockade acts to enhance CD28 signaling, for instance, which would result in activity in AP-1 heterodimers, c-Fos and c-Jun(*32*). This is notable given that exhausted T cells not only express high levels of inhibitory receptors but costimulatory receptors as well(*2*). One of the most well-known family members is the TNFR costimulatory receptor 4-1BB (CD137, encoded by *Tnfrsf9*), which is highly expressed on terminally exhausted T cells(*33*). T cell agonists, like agonistic antibodies to 4-1BB and OX40, are under intense clinical investigation, and we and others have previously shown that 4-1BB agonists could restore function in exhausted T cells(*34*). Indeed, 4-1BB signaling promotes activation of distinct AP-1 family members, such as ATF2, ATF3, and c-Jun(*35*).

We thus next asked whether checkpoint blockade or T cell agonist therapy could restore gene transcription at these anticorrelated loci. B16-bearing mice were treated with either anti-PD-1 or anti-4-1BB immunotherapy, and progenitor or terminally exhausted subsets were flow cytometrically sorted and sequenced by RNAseq (Fig 5A). Interestingly, only anti-4-1BB treatment led to increased expression of AP-1 family members including *Atf3*, *Batf3*, and *Mafb* (Fig 5B), yet both immunotherapies lead to enrichment in inflammatory response genes as well as cytokines and chemokines (Fig S5A,B). As expected, progenitor exhausted cells had no significant enrichment of anticorrelated genes between controls and treated animals, and limited enrichment of correlated genes, reflecting the limited expression of these gene sets in progenitor cells and both the likely limited targeting of cells with anti-4-1BB, and inability to capture progenitor cells post anti-PD-1 treatment (Fig S5C,D). Conversely, terminally exhausted cells isolated from anti-PD-1 treated mice were enriched in both correlated and anticorrelated genes, indicating that anti-PD-1 therapy leads to increased expression of genes with active chromatin states (Fig 5C, S5E). Strikingly, 4-1BB-treated exhausted T cells only increased expression of anticorrelative genes, with limited enrichment of correlated genes, suggesting that agonism of 4-1BB increases nuclear AP-1 and restores the expression of genes that retain active enhancers but lack gene expression at the steady state (Fig 5D, S5F).

**Figure 5.**
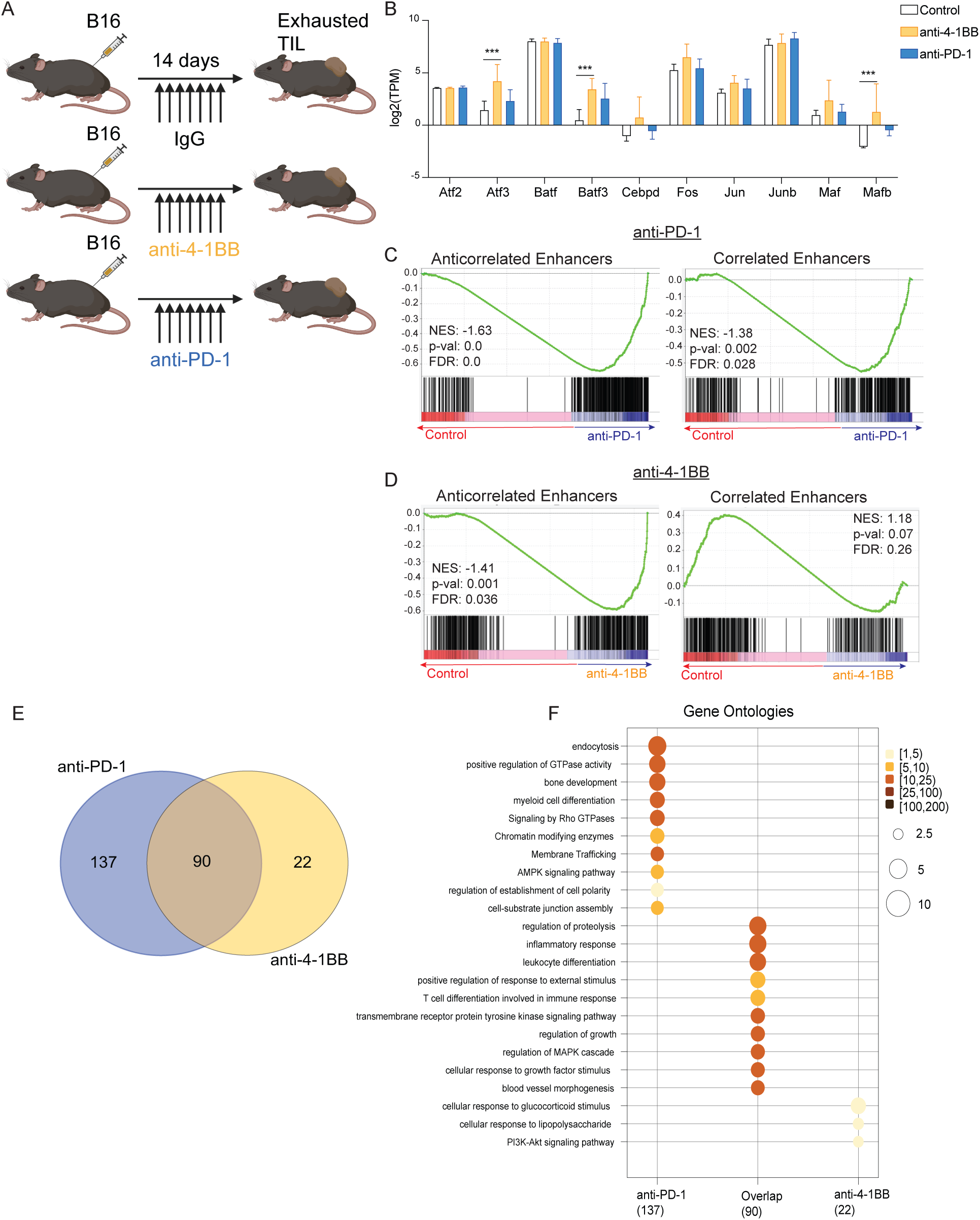
Immunotherapies that alter costimulatory signaling drive expression of anticorrelated genes. (A) Scheme of immunotherapy treatment of murine B16 tumors. (B) Log2 normalized expression (mean and SD) of AP-1 family members in terminally exhausted TIL in IgG-, anti-4-1BB-, or anti-PD-1-treated mice. p value (DESeq2, generated by one-way ANOVA); ***p<0.001 (C,D) Gene set enrichment analysis of transcriptomes of control (IgG) vs PD-1-treated (C) and 4-1BB-treated (D) mice. Gene lists of anticorrelated enhancers(left) and correlated enhancers (right) defined as in Figure 2 (Supplemental Table 1). (E) Venn diagram of terminal anticorrelated genes that change in expression upon treatment. (F) Bubble plot of enrichment of GO terms defined by Metascape in anticorrelated genes modified by treatment with anti-PD-1 or anti-4-1BB. RNAseq data was generated from three individual mice per treatment group sorted into progenitor (PD-1^lo^) or terminal (PD-1^hi^Tim3^+^) CD8 TIL.

As both anti-PD-1 and anti-4-1BB therapy increased expression of anticorrelated genes, we next asked if the genes restored by each therapy contained a core signature. Of the 249 restored genes, only 90 genes were commonly restored by both therapies while 137 genes or 22 genes were specifically changed by anti-PD-1 or anti-4-1BB alone respectively (Fig 5E). Pathway analysis of these 90 genes revealed significant enrichment in inflammatory response and leukocyte differentiation pathways, while genes impacted by anti-PD-1 alone were in endocytosis or GTPase pathways, and anti-4-1BB alone regulated responses to glucocorticoids (Fig 5F). Taken together, these data suggest that anti-4-1BB or anti-PD-1 therapy can restore expression of genes primed by an active chromatin landscape in terminally exhausted T cells, but only anti-4-1BB does so by increasing AP-1 family member expression, suggesting immunotherapies may be capable of “reinvigorating” some effector function of terminally exhausted cells by restoring gene expression of key inflammatory genes.

### Bivalent chromatin is increased in terminally exhausted TIL

In line with previous work, our data suggest that altering inhibitory receptor or costimulatory signaling via immunotherapy does not change the underlying epigenetics of exhausted T cells, but rather promotes transcription factor activity acting on the existing enhancer landscape(*8, 9*). Further, neither of these monotherapies can result in curative responses in B16 melanoma, suggesting other environmental factors may be at play that shape the epigenetics of T cells as they progress to exhaustion in cancer. Indeed, our data show that many regions of active chromatin, apart from these anticorrelated enhancers, were also present without corresponding gene expression, reminiscent of the poised state known as bivalency(*36*). Bivalent chromatin is defined as a state with both active H3K4me3 and repressive H3K27me3 with corresponding absence of gene expression(*37*). Increased states of chromatin bivalency have been described in stem cells and are thought to provide flexibility and readiness for subsequent differentiation upon appropriate extracellular signals. Given the increased state of active chromatin with anticorrelative gene expression in terminally exhausted T cells, we asked if there were intrinsic differences in bivalent chromatin in progenitor and terminal TIL. We used ChromHMM, a hidden Markov-model developed to characterize and identify patterns of chromatin marks to identify bivalent chromatin regions in TIL and OT-I CD8 effector T cells isolated from *Vaccina*-Ova (*38, 39*). To identify chromatin “states”, all TIL samples were pooled and states of chromatin were identified as permissive, bivalent, repressed or having no marks using both H3K4me3 and H3K27me3 (Fig 6A). IgG was included to eliminate any chromatin states that were only enriched as background. Three clusters were identified as repressed, based on the presence of only H3K27me3, where state 3 had that highest enrichment of H3K27me3. Two permissive states had enrichment of H3K4me3 alone with state 5 the predominant permissive state of the two. State 4 had similar enrichment profiles for both H3K4me3 and H3K27me3, indicating a chromatin state that was bivalent. Additionally, one state (state 7) had no enrichment of any marks. Seven chromatin states were also identified in OT-I effector cells, with a singular bivalent state present (Fig 6A). To determine if these defined states correlated with gene transcription, we assessed gene expression in terminally exhausted and OT-I effector cells using RNA-seq data to compare the mean transcripts present per chromatin state. Indeed, repressive states had on average reduced gene expression compared to permissive states (Fig 6B). Bivalent chromatin had reduced gene expression compared to permissive chromatin, in line with reports that genes with bivalent chromatin are not active(*36*) (Fig 6B). We next compared the frequency of chromatin states among the individual TIL populations in comparison to effector cells. While all TIL had increased levels of bivalent chromatin compared to effector cells, we were intrigued to find that the terminally exhausted cells had the largest percentage of bivalent chromatin (Fig 6C). This was unexpected as bivalency is normally associated with cellular flexibility and a stem cell like state rather than a terminal state of differentiation. Thus, these data suggested that increased bivalency may be another indication of a poised chromatin state leading to loss of gene expression in terminal T cell exhaustion.

**Figure 6.**
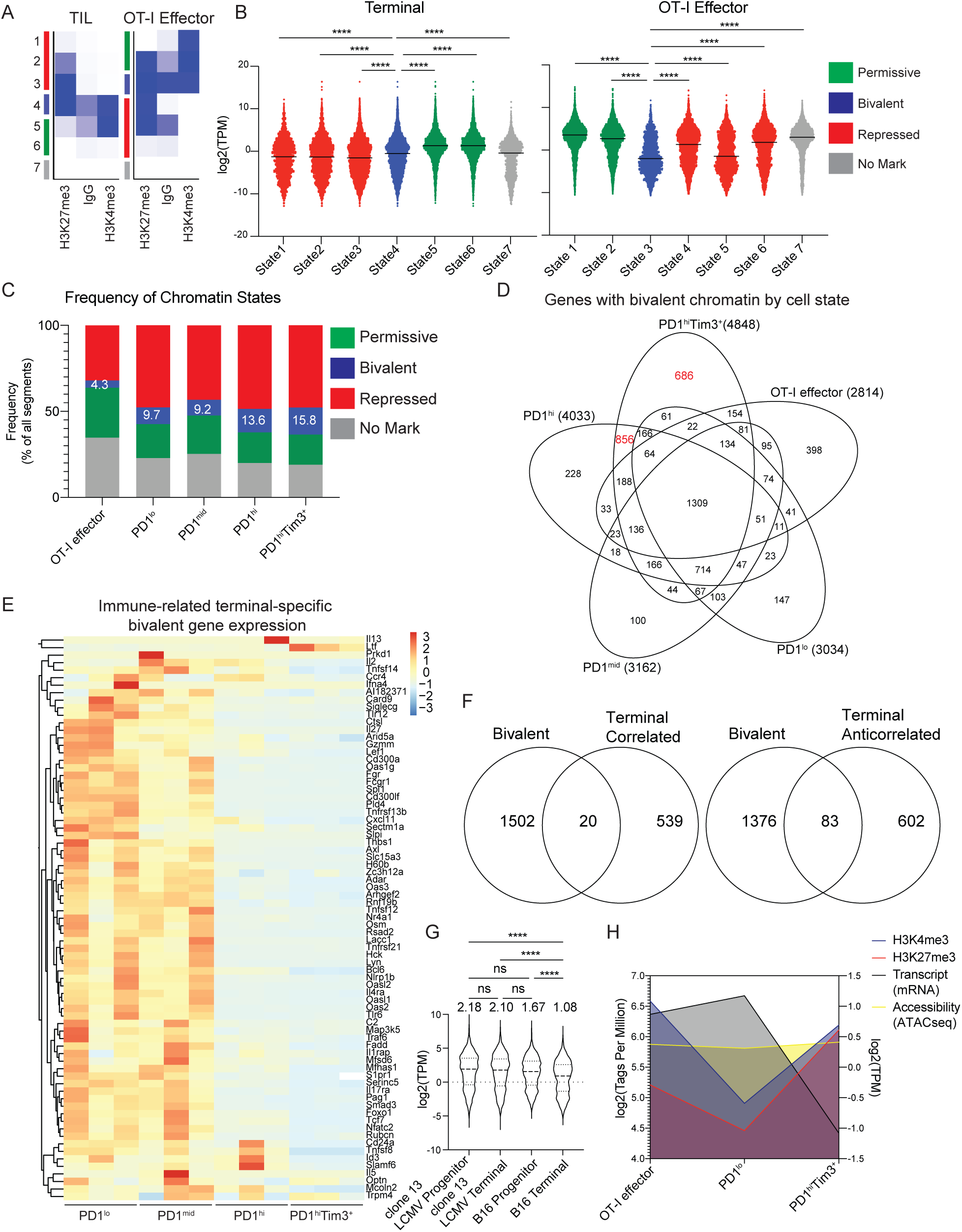
Terminally exhausted cells have increased bivalent genes, including important immune genes. (A) ChromHMM emissions plots depicting defined states in TIL and VVOVA effector cells. (B) Log2 normalized expression of genes nearest to the segments defined in A in each state in terminally exhausted TIL and VVOVA effector cells. (C) Frequency of chromatin states identified in A in TIL subsets and VVOVA effector cells. (D) Venn diagram of genes identified in each subset as bivalent based on the presence of bivalent chromatin defined in regions +/- 1kb of all TSSs and filtered for expression less than 2 TPM. (E) Heatmap of log2 normalized expression of select immune-related genes identified as exhaustion-specific bivalent genes. (F) Venn diagrams comparing exhaustion-specific bivalent genes to correlated and anticorrelated genes defined in Figure 2. (G) Violin plots of log2 normalized expression of bivalent genes identified in D from progenitor and terminally exhausted cells isolated from chronic LCMV infected mice or mice with B16 melanoma tumors (GSE122713). Numbers above each column indicate mean TPM value. One-way ANOVA, ****p<0.0001 (H) Summary plot showing changes in H3K4me3, H3K27me3, chromatin accessibility (ATAC-seq) and gene expression of exhaustion-specific bivalent genes identified in D. ATAC-seq (GSE122713) and *Listeria*-OVA infection (GSE95237).

We next explored the specific genes which exhibited bivalent chromatin in each TIL subset. We restricted our analysis to bivalent chromatin encompassing a 1kb region around the TSS and further removed any genes which had high gene expression, as this would not meet the criteria of bivalency. Indeed, terminally exhausted cells had an increased number of genes with bivalent promoter regions (unique to PD-1^hi^ Tim3^+^ (686) and shared with PD-1^hi^(856)) compared to effector (398) or progenitor exhausted cells (PD-1^lo^ (147); PD-1^mid^ (100))(Fig 6D). We found that terminal exhaustion bivalent genes were enriched for pathways involved in inflammatory response and leukocyte differentiation (Fig 6E, S6A,B). Importantly, the genes identified as bivalent have little overlap with the genes described in terminally exhausted cells as having active chromatin but anticorrelated gene expression, in line with comparable H3K27me3 at these regions (Fig 6F, Fig S3J). To determine if the terminal specific bivalent genes identified were also present in chronic viral infection, we explored their gene expression in previously published datasets of progenitor and terminal exhausted states isolated from chronic viral infection or B16 tumors(*6*). Gene expression was significantly decreased in tumor terminal exhaustion, but no change was observed in viral infection induced exhaustion, suggesting that the tumor microenvironment enforces a distinct terminal state in T cells and that the increase in bivalent chromatin is specific to tumor-mediated terminal exhaustion (Fig 6G).

Gene repression can be a function of chromatin accessibility. Thus, we explored the relationship between open or closed chromatin via ATACseq and bivalency. We assessed the amount of open chromatin, H3K4me3 and K3K27me3 via tag counts per peak together with corresponding transcript levels in OT-I effector cells from *Vaccinia*-Ova infection, progenitor and terminal exhausted TIL. We found no changes in chromatin accessibility at terminal-specific bivalent regions in effector, progenitor, or terminal cells despite changes in H3K27me3 and transcript expression (Fig 6H, S6C). Taken together, tumor-mediated terminal exhausted cells have an increase in bivalent chromatin at promoters leading to decreased gene expression.

### Tumor hypoxia leads to decreased gene expression at bivalent promoters in terminally exhausted cells

We recently reported that terminally exhausted cells carry severe metabolic defects, and part of the biology of exhausted T cell differentiation is the experience of oxidative stress and elevated reactive oxygen species (ROS)(*40*). Indeed, persistent antigenic stimulation under hypoxic conditions was sufficient to drive an exhausted-like fate *in vitro*. As oxygen is required for the dioxygenase reactions associated with histone and DNA demethylation, and terminally exhausted cells experience comparatively elevated hypoxia to progenitors, we asked whether environmental stressors like hypoxia can drive a bivalent state through alteration of demethylase activity(*41, 42*).

In T cells, the histone lysine demethylases (KDMs) that remove H3K27me3 are UTX and Jmjd3, encoded by *Kdm6a* and *Kdm6b* respectively(*43*). Numerous demethylases are downregulated in terminal cells, including both *Kdm6a* and *Kdm6b*, suggesting that H3K27 may not be sufficiently demethylated in terminal cells, potentially due to a combination of increased oxidative stress and reduced expression of KDMs (Fig 7A, Fig S7A). In contrast, *Ezh2*, which is responsible for methylating H3K27, was increased in terminal cells, further supporting a dysregulation in the balance of chromatin mediators that control methylation of H3K27(*44*)(Fig 7A).

**Figure 7.**
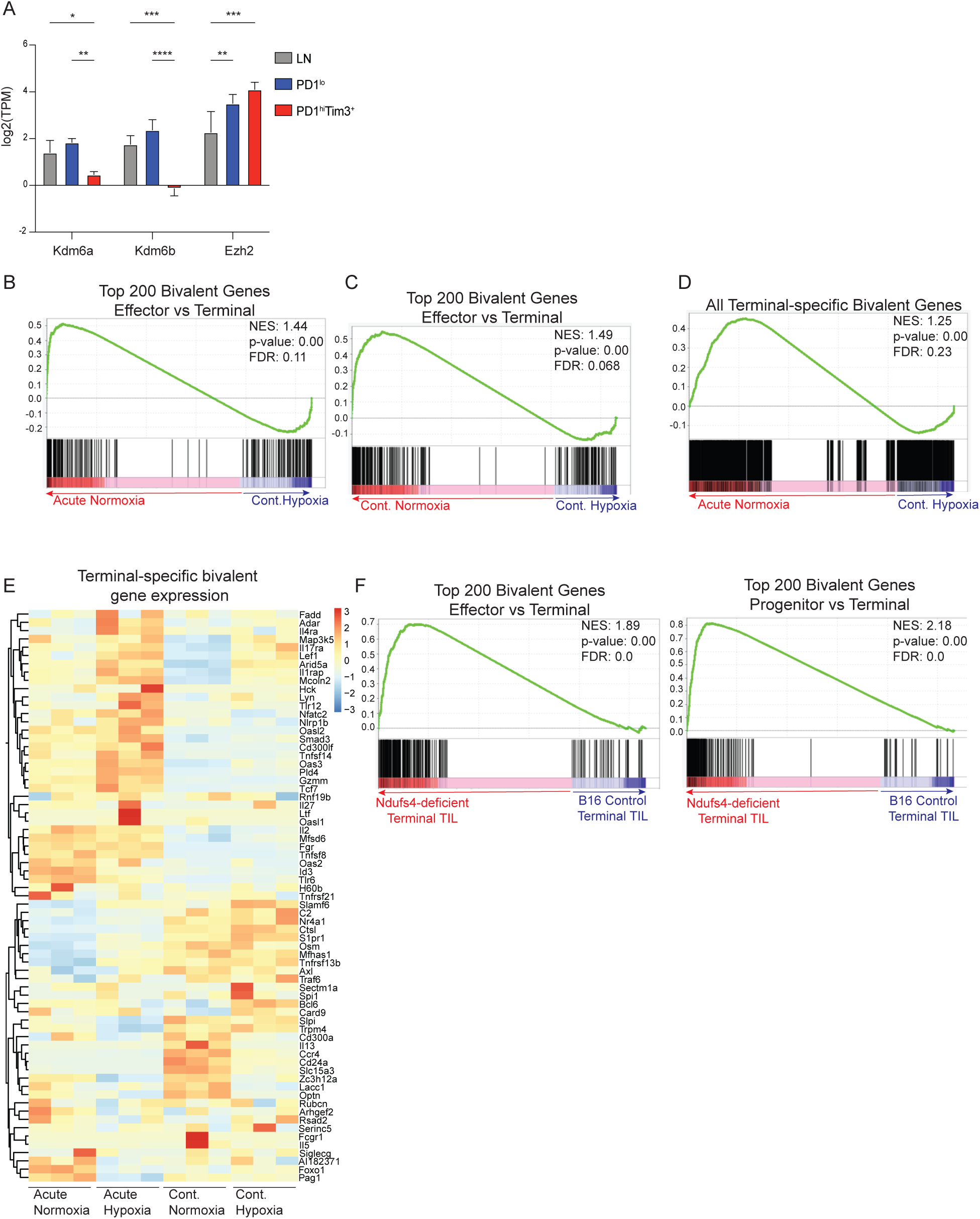
Hypoxia is sufficient to control expression of terminal-specific bivalent genes. (A) Log2 normalized expression (mean and SD) of histone modifiers known to regulate H3K27me3. p value (DESeq2, generated by one-way ANOVA); *p<0.05, **p<0.01,***p<0.001,****p<0.0001. (B-D) Gene set enrichment analysis of transcriptomes isolated from cells stimulated in vitro under conditions that drive exhaustion. (B,C) Top 200 bivalent genes defined in Fig 6D that decrease in expression from effector to terminally exhausted cells. (D) All bivalent genes in terminally exhausted cells defined in Fig 6D. (E) Heatmap of log2 normalized expression of select immune-related genes identified as bivalent in terminal exhaustion in *in vitro* stimulated cells. (F) Gene set enrichment analysis of terminally exhausted TIL transcriptomes isolated from *Ndufs4*-deficient or WT B16 melanoma tumors. Top 200 bivalent genes defined in Fig 6D that decrease in expression from effector to terminally exhausted cells (left) or from progenitor to terminally exhausted cells (right).

Histone demethylases such as *Kdm6a* are oxygen sensors, and hypoxic conditions are known to inhibit KDM activity, either directly or through hypoxia-inducible factor (HIF) mediated mechanisms(*45*). Notably, ROS, which are elevated in exhausted T cells as a consequence of metabolic dysfunction, acts as a competitive inhibitor of O_2_ binding for demethylases(*40, 46*). Thus, we hypothesized that the increased hypoxia experienced by terminally exhausted cells may contribute to the observed increase in bivalency. To determine if hypoxia can limit expression of genes identified as bivalent in terminally exhausted cells, we utilized an in vitro system of generating exhaustion-like cells by continuously stimulating CD8 T cells in the presence of hypoxia as we have previously reported(*40*). Unlike TIL cells, in vitro stimulation of cells in the presence of hypoxia did not alter expression of *Ezh2*, *Kdm6a* or *Kdm6b* (Fig S7B). Despite fewer changes in the expression of histone modifiers, we found that cells cultured with continuous stimulation in the setting of hypoxia had decreased expression of the most differentially regulated poised genes in terminally exhausted cells compared to effector T cells (Fig 7B). Similar results were achieved when comparing cells continuously stimulated in normoxia vs hypoxia, suggesting continuous stimulation was not the main driver of this transcriptional change (Fig 7C). Importantly, there was no enrichment in cells stimulated acutely or continuously in normoxia (Fig S7C). To determine if hypoxia could impact expression of genes identified as bivalent in terminally exhausted cells, we compared the transcriptomes of CD8 T cells cultured with acute stimulation in normoxia or hypoxia. Indeed, hypoxia downregulated many of the terminal-specific bivalent genes (Fig 7D), several of which are immune-related (Fig 7E). Taken together, these data suggested hypoxia is sufficient to drive downregulation of genes identified as bivalent in terminally exhausted cells.

To further delineate the relationship of hypoxia and bivalency and to determine whether reducing tumor hypoxia could reverse bivalency-driven gene repression, we utilized a genetically modified B16 tumor cell line engineered by CRISPR-Cas9 directed deletion of a key subunit of mitochondrial complex I (*Ndufs4*), resulting in a tumor that consumes less oxygen and produces less hypoxia but maintains baseline proliferation(*40*). In this context, targeting tumor hypoxia *in vivo* was sufficient to restore expression of Kdm6b (Fig S7D). Further, transcriptomic analysis of terminally exhausted (PD-1^hi^ Tim-3^+^) CD8 T cells sorted directly from Ndufs4-deficient B16 tumors were enriched for bivalent genes in comparison to terminal cells in control B16 tumors (Fig 7F, S7D). Thus, reducing hypoxia in the tumor increased expression of bivalent genes normally reduced in terminally exhausted T cells, suggesting hypoxia was responsible for driving bivalency and reducing gene expression. These data support a model in which the hypoxic environment of solid tumors drive epigenetic changes that underlie the exhaustion phenotype. Furthermore, reducing hypoxia in the tumor was sufficient to restore gene expression of bivalent genes, indicating that the epigenome of terminally exhausted CD8 T cells is not fixed but remains poised to re-express genes critical for effector functions. Our analysis of exhausted CD8 TIL has identified an epigenetic state in terminal exhaustion associated with a loss of gene expression that can be reacquired in certain settings. Thus, under the appropriate conditions anti-tumor immunity could be restored in terminally exhausted cells.

## DISCUSSION

Immunotherapeutic strategies targeting subsets of exhausted TIL have been limited by a lack of knowledge of the molecular details that underlie the transition from progenitor-like T cells (carrying potential to respond robustly to immunotherapy) to terminal exhaustion (which respond poorly to checkpoint blockade). How the tumor microenvironment impacts this progression at the epigenetic level has not previously been explored, and whether terminally exhausted T cells have therapeutic potential to gain effector capacity is unclear. We set out to profile the epigenetic landscape of CD8 TIL as they progressed from progenitor to terminally exhausted cells and compare them to bone fide effector and memory states. Our study has clarified the epigenetic landscape that underlies the transcriptional transition to terminal exhaustion and identified key epigenetic features that contribute to this state, as well as potential signals that drive them. By profiling four histone modifications that indicate active or repressed states of chromatin and the transcriptome of CD8 TIL as they progressed through exhaustion, we uncovered chromatin states with transcriptional potential that were limited by signals specific to the tumor microenvironment.

Active chromatin landscapes are marked by specific histone modifications that have been associated with transcription(*15*). When exploring regions of active chromatin that were unique to terminally exhausted cells, we were surprised to find a large percentage of genes with active chromatin landscapes yet reduced gene expression in terminal cells compared to progenitors. In contrast, progenitor and effector cells have active chromatin with mostly increased gene expression, suggesting these anticorrelated genes were specific to terminal exhaustion. Identifying with confidence specific enhancer-promoter relationships require chromatin conformation assays such as Hi-C, technologies that are currently limited by cell number or by the resolution of the chromatin space for closely related enhancer-promoter contacts. Thus, mapping or predicting enhancer-promoter contacts is an area of active research with no clear options given the low cell numbers of CD8 TIL isolated from murine tumors. Therefore, while distal enhancers could account for this lack of correlation, it would be surprising that terminally exhausted cells would have considerably more distal enhancer contacts than progenitor exhausted cells, suggesting the more likely hypothesis that there is a loss of enhancer-promoter contacts as cells progress to exhaustion. Active chromatin in terminally exhausted T cells was highly enriched for bZIP/AP-1 family motifs, yet very few AP-1 family members were expressed. Previous studies have shown that the AP-1 family plays a key role in selecting cell type-specific enhancer usage in macrophages and fibroblasts(*47, 48*). Thus, loss of AP-1 expression may be sufficient to limit enhancer-promoter contacts leading to loss of gene expression. Indeed, NFAT rendered incapable of binding AP-1 leads to increased exhaustion; thus, the balance of NFAT downstream of TCR and AP-1 downstream of co-stimulation plays a key part in the progression to exhaustion(*25, 31*). The precise timing and nature of the balance between TCR, co-stimulation and inhibitory receptor expression varies depending on the precise affinity of TCR interaction in the tumor, yet by the time cells exhibit features of terminal exhaustion they have lost AP-1 expression and subsequent gene expression. We hypothesize that sustained co-stimulation and AP-1 expression drive productive enhancer-looping to promote gene expression; however, more studies are needed to formally demonstrate the link between AP-1 expression and chromatin looping in CD8 TIL.

Immunotherapies that either target inhibitory receptor expression and directly agonize costimulatory molecules are known to alter signaling such as AP-1; thus, we reasoned that these strategies could restore expression of genes primed with active chromatin landscapes. We found that both PD-1 blockade and 4-1BB agonist treatments restored gene expression in terminal cells with active chromatin. However, 4-1BB specifically altered anticorrelative genes, while PD-1 blockade increased all gene expression in terminal cells with active chromatin. One challenge in comparing these two treatment strategies is the cellular targets are not similar. PD-1 primarily targets progenitor cells and allows progression to a more terminal state while effector function is maintained, whereas 4-1BB agonist acts on terminally exhausted T cells that exhibit elevated 4-1BB expression(*34*). We sorted progenitor and terminally exhausted cells separately to account for immunotherapeutic changes to the phenotype of the cells in bulk, but we are not able to parse these differences in a refined manner. While both treatments impacted anticorrelative gene expression, the precise genes within that group were not identical, suggesting PD-1 and 4-1BB therapy have different molecular signaling pathways that lead to increased effector cell gene programs. Yet both treatments were clearly able to restore gene expression, suggesting the balance of inhibitory and co-stimulatory signaling plays an important role in driving gene expression at regions with active chromatin.

While Tox is important for the transition to terminally exhausted T cells, Tox has other roles outside of exhaustion (*11, 12, 28*). Our analysis of Tox binding suggests it has roles in both progenitor and terminally exhausted states in the tumor. Motif analysis indicates that Tox pairs with distinct binding partners in each state, TCF-1 in progenitor and Batf:IRF (AICE elements) in terminally exhausted cells(*49*). TCF-1 can act as a pioneer factor, binding to regions of heterochromatin to promote opening and increased gene expression (*50*). However, intrinsic HDAC activity and repressive function has also been associated with TCF-1, suggesting that either the binding partners or the chromatin landscape impacts the precise role of TCF-1 in the context of exhaustion (*51*). Both Batf and Irf4 have been identified as key transcription factors regulating exhaustion(*14, 52, 53*). Complexes of Batf and IRF4 bind to AP-1:IRF composite elements (AICE) to drive differentiation of Th17 cells, B cells and dendritic cells(*49*). Intriguingly, Taiji analysis for terminal exhaustion ranked both Batf and Irf4 as important and positively correlated with expression, unlike AP-1 family members. Our data implicate a complex of Batf, Irf4, and Tox that is responsible for regulating gene expression in terminally exhausted cells. Tox is an HMG protein thought to bind DNA in a sequence-independent manner and has been associated with regulating chromatin architecture, suggesting that Tox may play a critical role in regulating enhancer-promoter chromatin looping (*27*).

Bivalent chromatin was initially described in pluripotent cells and is associated with genes that regulate cell fate decisions(*36*). The presence of both positive (H3K4me3) and negative (H3K27me3) chromatin states at a promoter region leads to a poised state, where gene expression can be driven by removal of H3K27me3, or more permanently repressed by removal of H3K4me3 leading to a heterochromatin state. Although technically challenging to demonstrate that bivalency exists in a single-cell at each allele for any given gene, recent advances in fluorescent molecule tracking and single cell genomic analysis suggest this state does exist at single nucleotide resolution(*54, 55*). Despite its association with a more flexible cell state, we identified an increase in bivalent chromatin in terminally exhausted cells. We hypothesized this was not due to a set of poised genes ready for further differentiation but rather was an aberration of lost demethylase activity. H3K27me3 is removed by two histone demethylases, *Kdm6a* (UTX) or *Kdm6b*(Jmjd3)(*56*). We found limited expression of both genes in terminally exhausted cells, while *Ezh2* expression, the methyltransferase responsible for adding methyl groups to H3K27 was intact. Hypoxia has been associated with increased bivalency, and we recently demonstrated terminally exhausted cells experience the most hypoxia, greatly impacting their function(*40, 45, 57*). Loss of function could be in part due to the ability of *Kdm6a* in particular to sense oxygen directly, which limits its function in low oxygen settings(*41, 42*). We found that changing hypoxia both *in vitro* and *in vivo* could alter expression of bivalent genes, supporting the notion that tumor hypoxia limits function of histone methylases and increases bivalency inhibiting gene expression. Importantly, the increased bivalency we identified is limited to specific genes and is not globally increased, suggesting a level of gene specific regulation must be occurring (data not shown). As there is interest in combining therapies targeting the epigenome including *Dnmt3a* and *Ezh2*, in combination with immunotherapies such as anti-PD-1 therapy to re-invigorate terminally exhausted T cells, further exploration of the molecular players driving epigenetic alterations such as bivalency require further study (*58, 59*).

Here, we have profiled the epigenome and associated transcriptome of CD8 TIL during the progression to exhaustion and described two unexpected chromatin signatures: the presence of active histone modifications without gene expression, and increased bivalency. Both lead to a loss in gene expression that appears primed for re-induction under settings that alter co-stimulatory signaling or tumor hypoxia. Our data highlight that it is a convergence of both altered immunologic signals and pathologic environmental signals that redirect differentiation to exhaustion and suggest that exhausted T cells remain in their dysfunctional state due to the memory of these previous stressors. However, our data collectively reveal a primed chromatin state in terminally exhausted T cells that may be available for reinvigoration given the right therapeutic inputs, and our study may be of use to develop new combination therapies that take full advantage of all subsets of tumor-infiltrating T cells to eradicate cancer cells.

## MATERIALS AND METHODS

### Design

Study design was a combination of controlled laboratory experiments and observational. Sample sizes for collection of sequencing analysis were determined based on cell numbers needed for sufficient technical quantities, and then repeated 2-3 times to generate biological replicates. All data was included in analysis and outliers only removed if there was sufficient evidence of contamination or technical flaws in the experimental process. Endpoints for all tumor experiments were in accordance with IACUC guidelines. Our research objective was to investigate the epigenetic and transcriptomic landscape of tumor-derived CD8 T cells and we used these datasets to form hypothesis such as the relationship of hypoxia and costimulatory signaling upon analysis of initial data.

### Mice

Animal work was done in accordance with the Institutional Animal Care and Use Committee of the University of Pittsburgh. All mice were on a C57BL/6 background, used at age 6–8 weeks, were both male and female, and housed in specific pathogen-free conditions before use. C57BL/6, SJ/L (Thy1.1), Tg(TcraTcrb)1100Mjb/J (OT-I) mice were obtained from the Jackson Laboratory.

### Cell lines and Virus

B16-F10 cells were obtained from the American Type Culture Collection. *Ndufs4*-deficient B16-F10 mice were generated using transient transfection of Cas9–GFP and *Ndufs4*-targeted guide RNA, single-cell sorting of GFP^+^ cells, *in vitro* culture and screening using extracellular flux analysis, and selection of a clone that had lost GFP expression and lacked Ndufs4 protein. *Vaccinia*-OVA was generated by J. R. Bennink and provided by J. Powell (Johns Hopkins University). Zombie viability dye and anti-CD8 (53.6.7), anti-Tim-3 (RMT3-23), anti-PD-1 (29F.1A12), anti-CD44 (IM7), anti-Thy1.1 (OX-7), anti-CD105 (MJ7/18) antibodies, and Mojo Buffer and MojoSort Streptavidin Nanobeads were obtained from BioLegend. Dynabeads Mouse T-Activator CD3/CD28 for T cell expansion and activation were obtained from Fisher. IL-2 and IL-12 were obtained from PeproTech*. In vivo* antibodies anti-PD-1 (clone J43), anti-4-1BB (clone 3H3) and isotype controls were obtained from Bio X Cell.

### Tumor Experiments

Mice were injected intradermally with 2.5 × 10^5^ B16 melanoma cells. Mice were sacrificed and dLN and TIL harvested on day 14, when tumors were typically 8-10 mm in diameter. In immunotherapy experiments, when tumors were palpable (typically day 5), mice began anti-PD-1, anti-4-1BB, or isotype control therapy. Mice were treated with 200 μg of anti-PD-1, 200 μg of anti-4-1BB, or equivalent isotype control intraperitoneally three times per week for the duration of the experiment.

### *Vaccinia*-OVA Experiments

Mice were simultaneously injected with 10^6^ PFU of *Vaccinia*-OVA intraperitoneally and 1 × 10^5^ naïve OT-1^+^Thy1.1^+^CD8^+^ T cells retro-orbitally. On day 10, spleens were harvested, and erythrocytes lysed with a 1x solution of RBC lysis buffer (Thermo Fisher). Thy1.1^+^ T cells were enriched by negative magnetic bead selection using MojoSort Streptavidin Nanobeads (Biolegend) and biotinylated antibodies: anti-TCRγ/δ, anti-CD19, anti-B220, anti-NK1.1, anti-CD49b, anti-CD105, anti-CD32/64, anti-CD11c, anti-CD11b, anti-Ly6G, and anti-CD24. Activation of OT-1^+^CD8^+^ T cells was verified by flow cytometry.

### T Cell Isolation

For TIL isolation, surgically excised tumors were injected repeatedly with a total of 2 mL of 2 mg/ml collagenase type IV, 2 U/ml of hyaluronidase, and 10 U/ml DNase I in RPMI with 10% FBS and incubated for 30 minutes at 37°C. Tumors were mechanically disrupted between frosted glass slides, filtered, and vortexed for 30 seconds. Immune cell populations were ‘tumor debulked’ by negative magnetic bead selection (MojoSort; Biolegend) using anti-CD105. Lymph node and splenic T cells were isolated by mechanic disruption between frosted glass slides and filtered. Erythrocytes were then lysed as before.

### Cell Staining and Sorting

After lymph node and TIL were dissociated and enriched by MojoSort negative selection, preps were stained in 1x PBS with Zombie viability dye, anti-CD8, anti-Tim-3, anti-PD-1, and anti-CD44 for 15 minutes on ice. Following wash cycles by centrifugation in 1x PBS, cells were resuspended at 2 × 10^7^ cells per mL in sterile sorting buffer (500 mL 1x PBS, 2.5 g BSA, 12.5 mL 1 M HEPES, 5 mL 500 nM EDTA) and maintained on ice and shielded from light before fluorescence activated cell sorting via Beckman Coulter MoFlo Astrios or Sony MA900.

### *In vitro* T cell culture

For the bead-based continuous stimulation under hypoxia assay, LN and spleen CD8^+^ T cells were isolated from 6- to 8-week-old mice, mechanically disrupted and sorted on CD8^+^CD44^hi^ via Beckman Coulter MoFlo Astrios or Sony MA900. Cells were then activated at 2 × 10^4^ T cells per well in 96-well round-bottomed plates with an equivalent number of CD3/CD28 washed dynabeads (Thermo Fischer), 25 U ml^−1^ of IL-2, and 10 ng ml^−1^ of IL-12 in 200 μl of complete RPMI + 10% serum (starting point = day 0). Cells were activated for 24 h, then the beads were magnetically removed, and the cells divided into four conditions: no dynabeads in regular incubator (acute activation in normoxia), no dynabeads in 1.5% oxygen hypoxia chamber (acute activation in hypoxia; BioSpherix, ProOx Model C21), and with 200,000 dynabeads (1 cell per 10 dynabeads) in the regular incubator (continuous activation in normoxia), and with 200,000 dynabeads in a 1.5% oxygen hypoxia chamber (continuous activation in hypoxia). Cells were cultured in 25 U ml^−1^ of IL-2 in 300 μl of complete RPMI + 10% serum in 96-well round-bottomed plates (starting culture conditions = day 1). After 48 h (day 3), cells were split in half (for example, 10 wells per group are now 20 wells per group), with fresh medium + IL-2 used to replace the old medium. Bead numbers were kept consistent per well. After 48 h (day 5), cells were split in half again, with fresh medium + IL-2 used to replace the old medium. Bead numbers were kept consistent per well. After 24 h (day 6), cells were assayed after the beads had been removed.

### RNA Sequencing

cDNA was prepared from ∼1,000 cells using the SMART-seq v4 Ultra Low Input RNA Kit for Sequencing, (Clontech Laboratories). Sequencing libraries were prepared using the Nextera XT DNA Library Preparation kit (Illumina), normalized at 2nM using Tris-HCl (10 mM, pH 8.5) with 0.1% Tween-20, diluted and denatured to a final concentration of 1.8nM using the Illumina Denaturing and Diluting libraries for the NextSeq 500 protocol Revision D (Illumina). Cluster generation and 75-bp paired-end, dual-indexed sequencing were performed on the Illumina NextSeq 500 system.

### CUT&RUN Assay

CUT&RUN assay was performed as previously described (*17*). Live sorted cells were incubated overnight with Concanavalin A beads and antibodies recognizing H3K4me3(Abcam), H3K27me3 (Cell Signaling Technology), H3K27ac (Abcam), H3K9ac (Abcam), Tox (Abcam), Batf (Brookwood Biomedical), or IgG(Cell Signaling Technology). pA-MNase was then added, followed by CaCl_2_ to cleave the antibody-bound chromatin. Phenol-chloroform extraction was then performed to isolate enriched DNA. Two replicates were performed for each antibody (exception, Batf). Libraries were prepared using NEBNext Multiplex Oligos for Illumina and either the sparQ DNA Library Prep Kit (Quantabio) or NEBNext Ultra II DNA Library Prep Kit for Illumina. Libraries were quantified by qPCR using either the sparQ Library QuantKit (Quantabio) or NEBNext Library Quant Kit for Illumina. Appropriate library size was confirmed by running amplified qPCR products on an agarose gel. Cluster generation and 75-bp paired-end, dual-indexed sequencing were performed on the Illumina NextSeq 500 system.

### Sequencing Data Pre-Processing

FastQC was used to perform a quality assessment on all fastq files. The mouse reference genome (GRCm38) was downloaded from Ensembl. Adaptors were trimmed for both RNA sequencing and CUT&RUN data using cutadapt v1.18. RNA sequencing samples were aligned to using HISAT2 v2.1.0. Raw count values were generated using Subread, and gene expression values were normalized using transcripts per million. CUT&RUN reads were aligned to the reference genome using Bowtie2 v2.3.4.2. Duplicates were removed using Picard, and regions from the ENCODE Blacklist were removed using bedtools intersect.

### Peak Calling and Generation of Tag Count Files

For the characterization of reproducible histone modification and transcription factor peaks from CUT&RUN and publicly available ATAC-seq and ChIP-seq datasets, macs2 v2.1.1 was applied. For H3K27ac, a p value cutoff of 0.05 was used. For all other marks and ATAC-seq, a p value cutoff of 0.005 was used. Broad peaks were called for H3K27me3 and H3K4me3, whereas narrow peaks were called for H3K27ac, H3K9ac, ATAC-seq, and transcription factors. Irreproducible discovery rate (IDR) peaks were then found using a threshold of 0.05. IDR peak files of all TIL samples were then merged to create a master list of peaks for each mark. The master peak list and the alignment files were then used in bedtools coverage to generate tag counts for each replicate.

### Differential Peak and Gene Expression Analysis

DESeq2 was applied to raw counts transcriptome data in a pairwise manner to determine differentially expressed genes. These gene lists were compiled to make a master list of all differentially expressed genes. Log2 transformed normalized counts were then displayed as a heatmap to display expression of differentially expressed genes. DESeq2 was applied to raw tag count data for called peaks in a pairwise manner to determine differential peaks. These peak lists were compiled to make a master list of all differential peaks for each mark. Peaks were annotated to the nearest gene using ChIPpeakAnno (Zhu et al 2010).

### Analysis of Chromatin Data

Motif enrichment analysis was performed using HOMER findMotifsGenome (Heinz et al, 2010). Differential peak files were used for these analyses to define the specific motifs that were changing. The background file was specified using progenitor as background for terminal analyses and vice versa, unless otherwise specified. All motifs shown are from the known motifs output of findMotifsGenome.

HOMER makeTagDirectory and AnnotatePeaks functions were used to make histograms. Alignment files were used in makeTagDirectories to make tag directories for each replicate. AnnotatePeaks was used with the peak file of interest and the tag directories to generate magnitude data for each replicate at a given point from the center of the peak. These values were normalized using the read count for each replicate and graphed using Graphpad Prism.

PageRank analysis was performed using Taiji(*26*).The analysis was performed using H3K4me3, transcriptome, and H3K27ac data to rank the transcription factors. The motifs used were from the Homer database. Fold change between progenitor and terminally exhausted PageRank values was then calculated to determine factors important in each context.

Bivalent chromatin was defined using ChromHMM. Bam files with duplicated and blacklisted regions removed were used in the BinarizeBam function with a bin size of 1000. The LearnModel function was then used with 7 states defined and a bin size of 1000. Bivalent states were defined using the presence of both H3K27me3 and H3K4me3 and low gene expression.

## Data and materials availability

The datasets generated during this study are available at NCBI GEO repository GSE175443. Additional datasets not generated by this study are available at NCBI GEO repository GSE155192, GSE123235, GSE123236, GSE95237, GSE54191.

## Statistical Analysis

Statistical significance of genomics data was determined using p values given by DESeq2. Any other methods of determining statistical significance are described in the figure legend.

## Acknowledgments

We are grateful to Dr. Steven Henikoff (Fred Hutch Cancer Center) for initial pA-MNase reagent and Dr. Sarah Hainer (U. Pittsburgh) for pA-MNase plasmid and associated CUT&RUN technical advice. We thank the UPMC Hillman Cancer Center Cytometry Flow core and the UPMC Children’s Hospital of Pittsburgh Flow Core for help with cell sorting and analysis. We also thank the Health Sciences Sequencing Core at UPMC Children’s Hospital of Pittsburgh for Next Generation Sequencing Services, and the University of Pittsburgh Center for Research Computing for computing cluster access and support.

## Funding

B.R.F. supported by T32 CA082084; N.E.S supported by the NCI Predoctoral to Postdoctoral Fellow Transition Award (F99/K00) (no. F99CA222711). P.D.A.V. supported by the National Cancer Institute of the National Institutes of Health (NIH) Ruth L. Kirschstein National Research Service Award 1F30CA247034-01 and T32CA082084, and the National Institute of General Medical Sciences of the National Institutes of Health (T32GM008208). R.P. was supported by T32AI089443. A.C.P. supported by R01AI153104, R21AI135027. G.M.D. was supported by an NIH New Innovator Award (DP2AI136598) and R21AI135367, the UPMC Hillman Cancer Center Melanoma/Skin Cancer (P50CA121973) and Head and Neck Cancer SPORE (P50CA097190), the Alliance for Cancer Gene Therapy, the Mark Foundation for Cancer Research Emerging Leader Award, the Cancer Research Institute Lloyd J. Old STAR Award, and the Sy Holzer Endowed Cancer Immunotherapy Fund. This work utilized flow cytometry and animal facilities at UPMC Hillman Cancer Center, supported by P30CA047904.The content is solely the responsibility of the authors and does not necessarily represent the official views of the National Institutes of Health.

## Author Contributions

Conceptualization: N.E.S., B.R.F., N.L.R., A.C.P., G.M.D.

Methodology: N.E.S., P.D.A.V., N.L.R., B.R.F., A.C.P., G.M.D.

Software: N.L.R. B.R.F.

Validation: N.L.R. B.R.F.

Formal Analysis: B.R.F., N.L.R., N.E.S., P.D.A.V., A.C.P.

Investigation: B.R.F., N.L.R., N.E.S., P.D.A.V., A.T.F., R.P.

Resources: A.C.P., G.M.D.

Data Curation: N.L.R., B.R.F.

Writing – Original Draft: B.R.F., A.C.P., G.M.D.

Writing – Review & Editing: N.E.S., P.D.A.V., N.L.R., B.R.F., A.C.P., G.M.D.

Visualization: B.R.F., N.L.R., N.E.S., P.D.A.V., A.C.P.

Supervision: A.C.P., G.M.D.

Project Administration: A.C.P., G.M.D.

Funding Acquisition: A.C.P., G.M.D.

## Competing Interests

B.R.F., P.D.A.V., N.L.R., N.E.S., A.T.F., R.P., A.C.P. have no interests to declare. G.M.D. declares competing financial interests and has submitted patents covering the use of metabolic reprogramming in cell therapies that are licensed or pending and is entitled to a share in net income generated from licensing of these patent rights for commercial development. G.M.D. consults for and/or is on the scientific advisory board of BlueSphere Bio, Century Therapeutics, Nanna Therapeutics, Novasenta, Pieris Pharmaceuticals, and Western Oncolytics/Kalivir; has grants from bluebird bio, Nanna Therapeutics, Novasenta, Pfizer, Pieris Pharmaceuticals, TCR2, and Western Oncolytics/Kalivir; G.M.D. owns stock in Novasenta.

---

**Supplementary Figure 1.**
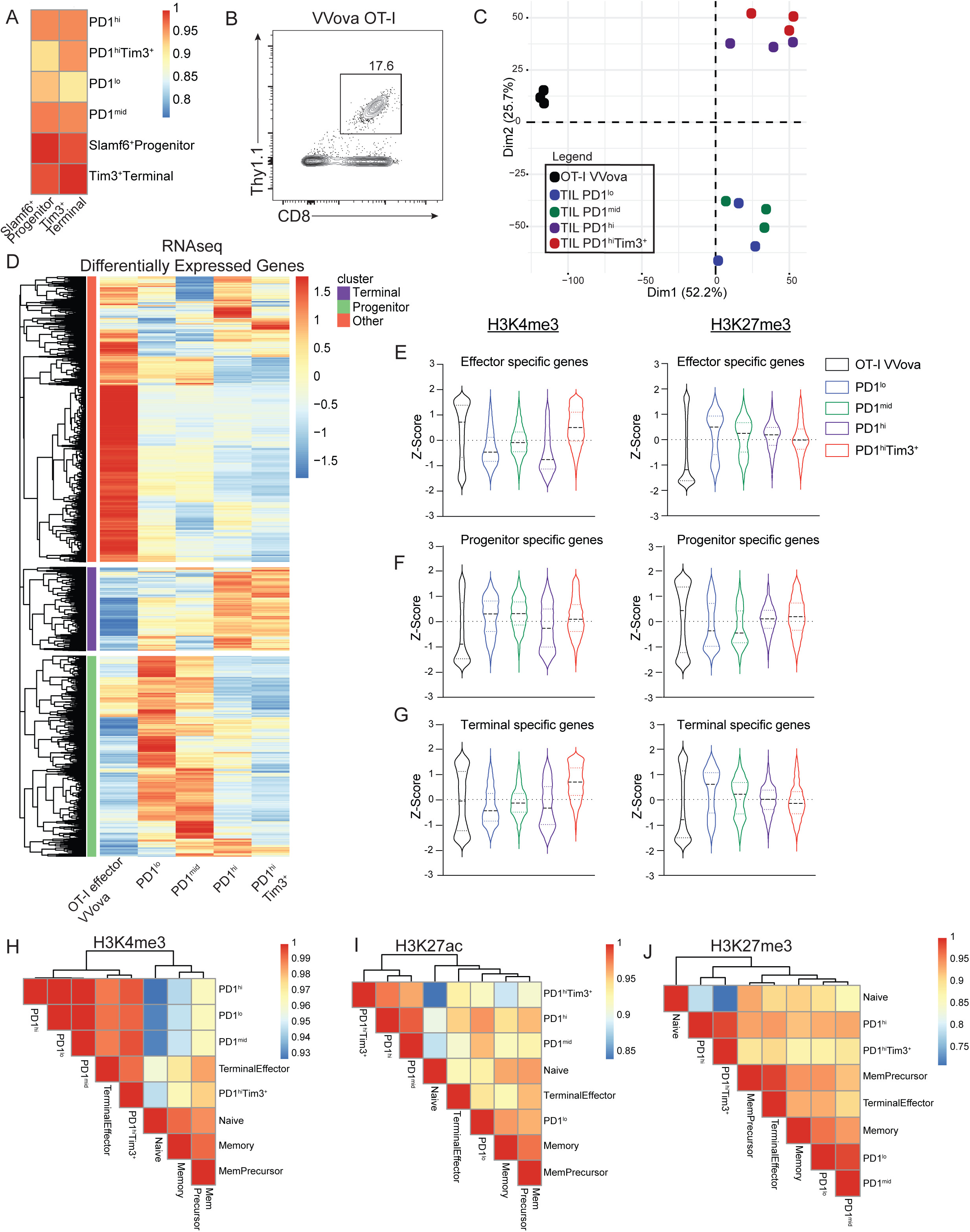
Exhausted tumor-infiltrating T cells are transcriptionally and epigenetically distinct from effector and memory T cells. (A) Pearson correlation showing relationship between RNAseq data for TIL subsets and RNAseq data for described progenitor and terminally exhausted subsets (GSE123235). (B) Thy1.1+ CD8 OT-I effector T cells were sorted from spleens from day 8 of VVOVA-infected C57Bl/6 mice. (C) PCA plot comparing transcriptome data from day 8 OT-I VVOVA effector T cells to TIL subsets from B16 melanoma. (D) Heatmap of log2 normalized expression of differentially expressed genes between day 8 VVOVA effector T cells and TIL subsets from B16. (E-G) Violin plots of H3K4me3 and H3K27me3 coverage of genes upregulated in effector (E), PD-1^lo^ (F), and PD-1^hi^Tim3^+^ (G) cells. Z-score indicates tag counts per million of 10kb regions surrounding the TSS. (H-J) Pearson correlation showing relationship between CUT&RUN of TIL subsets and publicly available ChIP-seq from naïve, terminal effector, memory progenitor, and memory T cells from Listeria-OVA infection for H3K4me3 (H), H3K27ac (I), and H3K27me3 (J) (GSE95237).

**Supplementary Figure 2.**
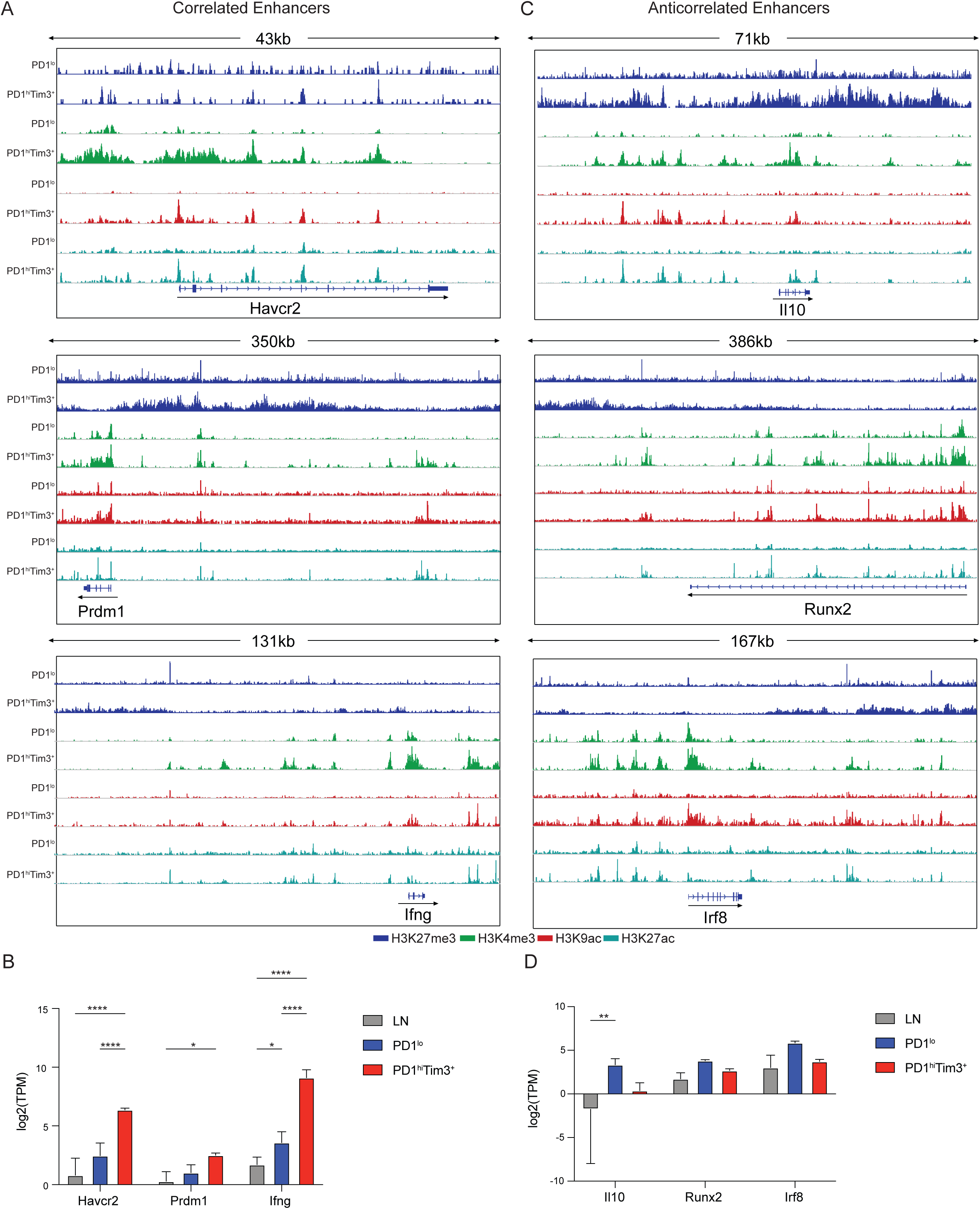
Examples of correlated and anticorrelated enhancers. (A) Representative IGV plots showing changes in H3K27me3, H3K4me3, H3K9ac, and H3K27ac at the correlated genes *Havcr2*, *Prdm1*, and *Ifng*. (B) Log2 normalized gene expression (mean and SD) of the correlated genes *Havcr2*, *Prdm1*, and *Ifng*. p value (DESeq2); *p<0.05, **p<0.01,***p<0.001,****p<0.0001. (C) IGV plots showing changes in H3K27me3, H3K4me3, H3K9ac, and H3K27ac at the anticorrelated genes *Il10*, *Runx2*, and *Irf8*. (D) Log2 normalized gene expression (mean and SD) of the correlated genes *Il10*, *Runx2*, and *Irf8*. p value (DESeq2); *p<0.05, **p<0.01,***p<0.001,****p<0.0001.

**Supplementary Figure 3.**
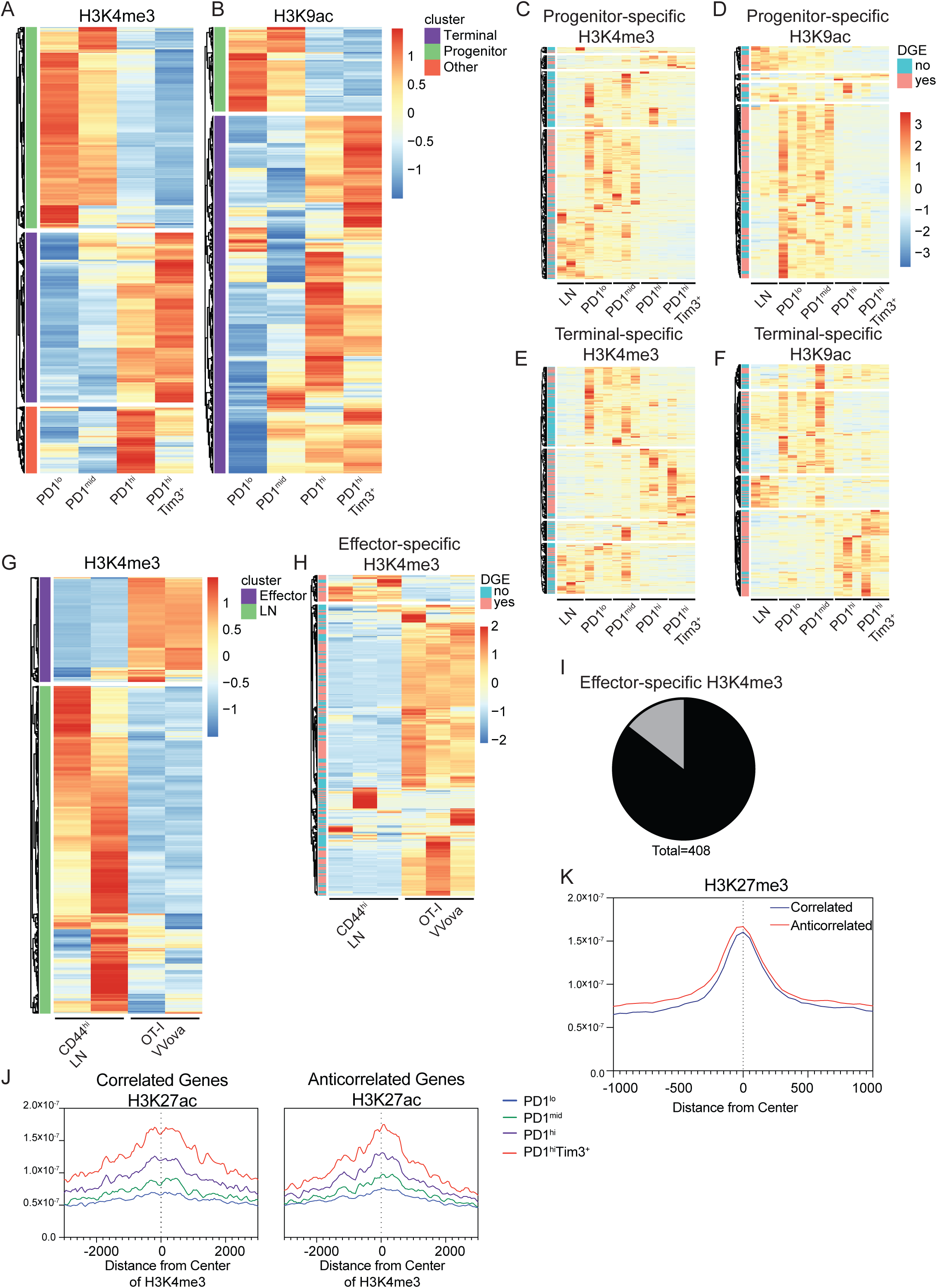
Anticorrelated and correlated genes have comparable levels of active and repressive marks. (A and B) Heatmap shows DESeq2 log2 normalized tag counts of (A) H3K4me3 or (B) H3K9ac at differential peaks identified between TIL subsets. (C-F) Heatmaps of log2 normalized expression of genes nearest to progenitor-specific (C) and terminal-specific (D) H3K4me3 peaks or progenitor-specific (D) and terminal-specific (F) H3K9ac peaks. Those genes that meet differentially expressed genes cutoffs (FC >2, p val < 0.05) marked in pink on the left side of heatmap. LN is CD44^hi^ CD8+ (G) Heatmap shows DESeq2 log2 normalized tag counts of H3K4me3 differential peaks between LN and VVOVA effector cells. (H) Heatmap of log2 normalized expression of genes nearest to effector-specific H3K4me3 peaks. Those genes that meet differentially expressed genes cutoffs (FC >2, p val < 0.05) marked in pink on the left side of heatmap. (I) Histograms showing H3K4me3 coverage at correlated and anticorrelated H3K27ac terminal exhaustion specific peaks. (J) Histogram showing H3K27me3 coverage at anticorrelated and correlated H3K27ac terminal exhaustion specific peaks.

**Supplementary Figure 4.**
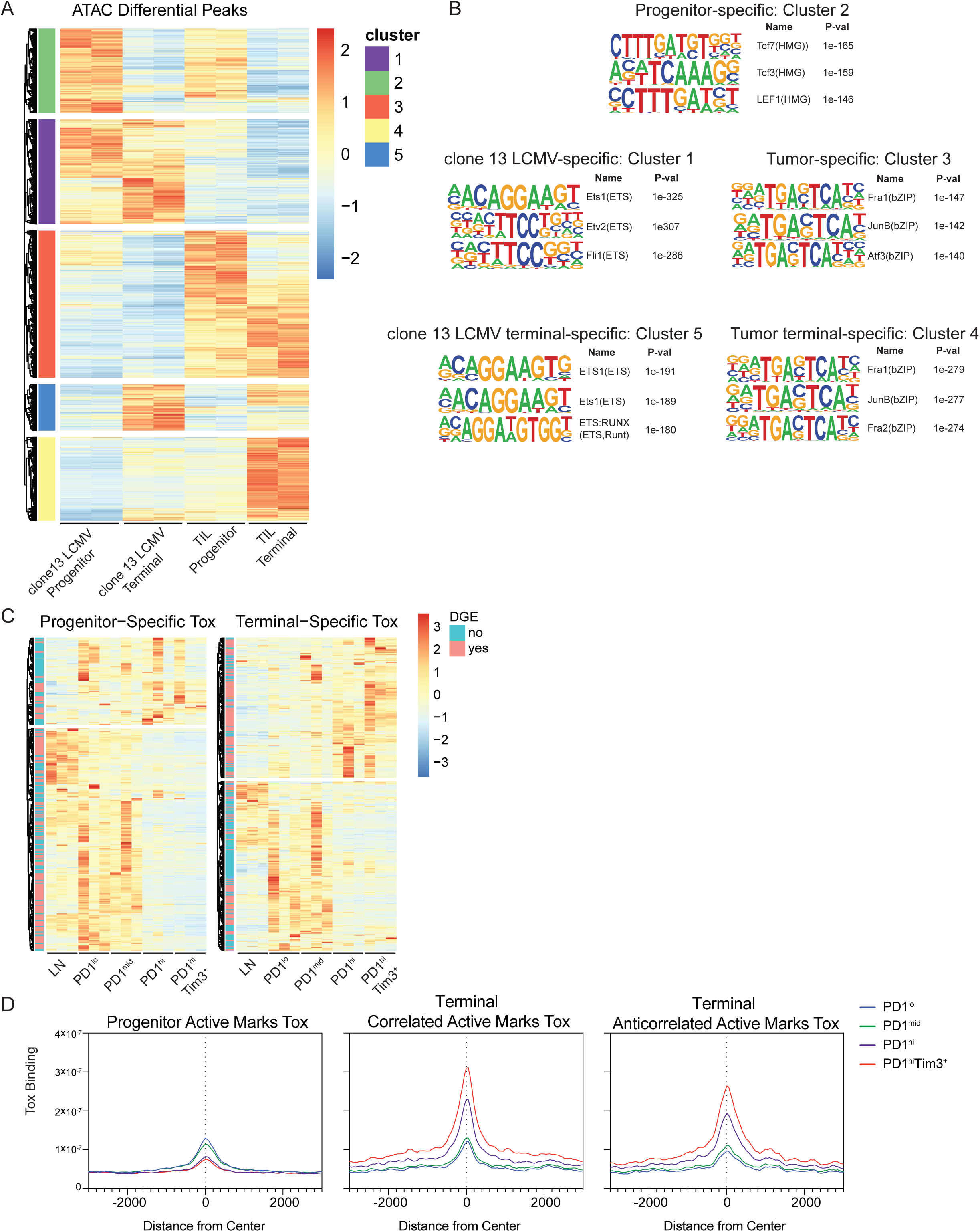
Tumor-specific bZIP enrichment and Tox binding at active chromatin. (A) Heatmap showing log2 normalized DESeq2 defined differential accessibility peaks between exhausted subsets from chronic LCMV and B16 melanoma(GSE123235). (B) HOMER motif analysis for each cluster defined in (A). Default background was used for all motif analyses. (C) Heatmaps showing log2 normalized expression of genes nearest to progenitor-specific and terminal-specific Tox peaks. Those genes that meet differentially expressed genes cutoffs (FC >2, p val < 0.05) marked in pink on the left side of heatmap. (D) Histograms showing Tox coverage at progenitor, correlated, and anticorrelated active mark peaks.

**Supplementary Figure 5.**
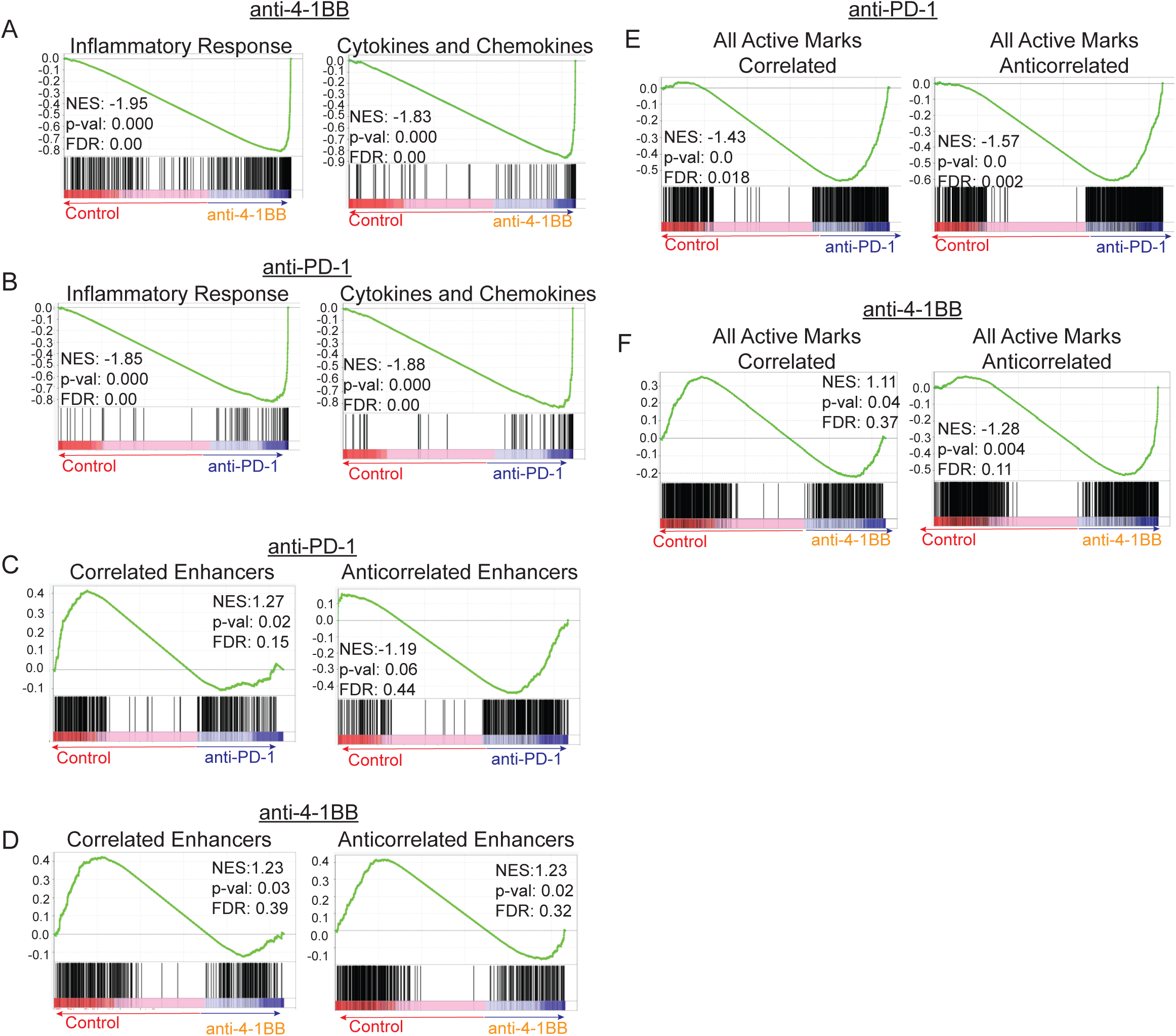
Immunotherapies have distinct effects on correlated and anticorrelated genes. (A and B) Gene set enrichment analysis for inflammatory response and cytokines and chemokines gene sets, comparing terminally exhausted TIL from control and 4-1BB-treated mice (A) or control and PD-1-treated mice (B). (C and D) Gene set enrichment analysis for correlated or anticorrelated enhancer gene sets, comparing terminally exhausted TIL from control and PD-1-treated mice (C) or control and 4-1BB-treated mice (D). (E and F) Gene set enrichment analysis for all active marks with correlated or anticorrelated gene expression, comparing terminally exhausted TIL from control and PD-1-treated mice (E) or control and 4-1BB-treated mice (F).

**Supplementary Figure 6.**
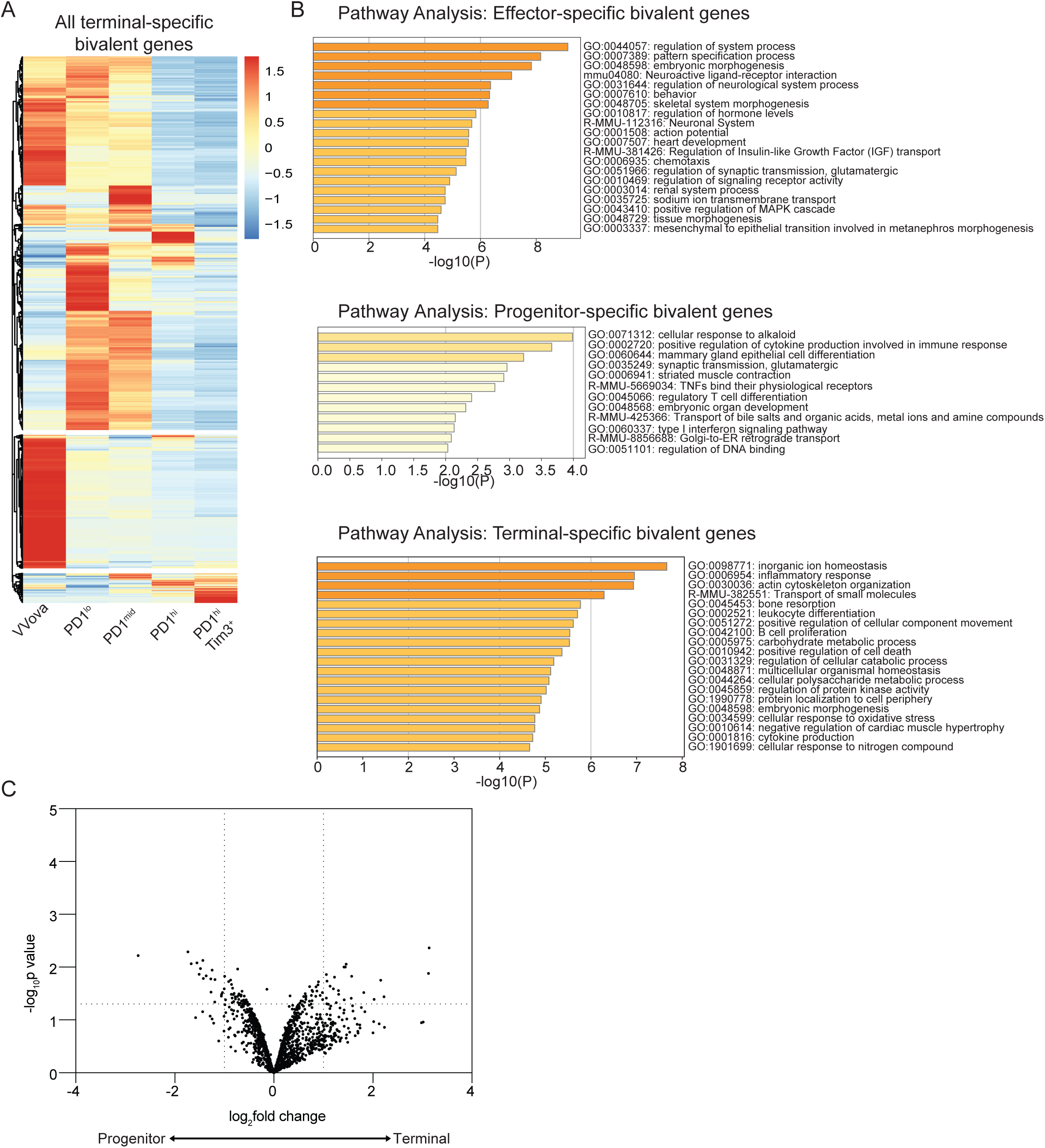
Terminal-specific bivalent genes have decreased expression but limited change in chromatin accessibility. (A) Heatmap of log2 normalized gene expression of terminal-specific bivalent genes in day 8 VVOVA effector T cells and TIL subsets from B16. (B) Enrichment of GO terms defined by Metascape in effector-, progenitor-, and terminal-specific bivalent genes. (C) Volcano plot of fold change and p value of changes in chromatin accessibility determined by ATAC-seq (GSE122713) in progenitor and terminally exhausted cells from B16 melanoma at terminal-specific bivalent genes (Log2 fold-change > 1 and log10 p value < 0.05). Coverage was determined for 10kb regions surrounding the TSS of each gene.

**Supplementary Figure 7.**
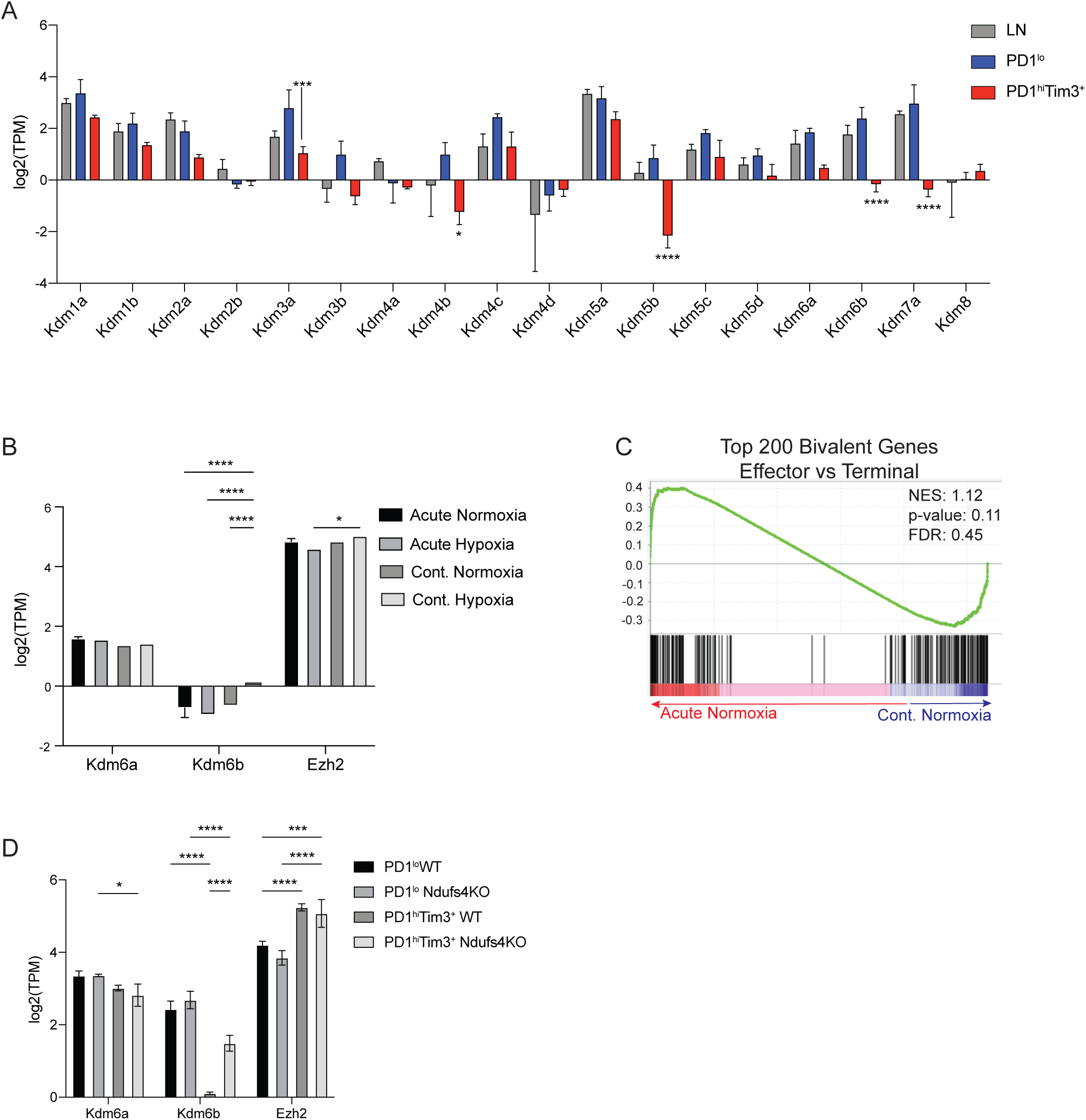
Hypoxia regulates expression of histone modifiers. (A) Log2 normalized gene expression (mean and SD) of histone demethylase genes in LN, PD-1^lo^, and PD-1^hi^Tim3^+^. p value (DESeq2); *p<0.05, **p<0.01,***p<0.001,****p<0.0001. (B) Log2 normalized gene expression (mean and SD) of histone modifiers that regulate H3K27me3 for acute and continuously stimulated cells in hypoxia or normoxia. p value (DESeq2); *p<0.05, **p<0.01,***p<0.001,****p<0.0001 (C) Gene set enrichment analysis for top 200 bivalent genes decreasing in expression from effector to terminally exhausted cells comparing acute stimulation in normoxia to continuous stimulation in normoxia. (D) Log2 normalized gene expression (mean and SD) of histone modifiers that regulate H3K27me3 for progenitor and terminally exhausted TIL from WT and *Ndufs4*-deficient B16 melanoma. p value (DESeq2); *p<0.05, **p<0.01,***p<0.001,****p<0.0001

**Supplemental Table 1: Gene lists for GSEA analysis**

Gene lists used in GSEA analysis were curated from datasets or from pathway focused gene expression lists.

